# REWRITE Enables Locus-Scale Engineering of Synthetic HLA Haplotypes in Human Pluripotent Stem Cells

**DOI:** 10.1101/2025.09.16.676382

**Authors:** Serena F. Generoso, Sarah Levovitz, Susanna Jaramillo, Minjoo Kim, Sumanth Dara, Shean Fu Phen, Bryan Yi, Tomoki Yanagi, Thomas L. DesMarais, Neta Agmon, Megan S. Hogan, Leslie A. Mitchell, David M. Truong

## Abstract

Human leukocyte antigen (HLA) polymorphism underlies antigen presentation, immune recognition, and central tolerance, yet the complex, multigenic structure of native HLA loci has precluded systematic engineering in human pluripotent stem cells (hPSCs). Existing genome engineering demonstrations in hPSCs focus on smaller integrations or at ectopic sites, leaving scar-minimized rewriting of native human loci at >100 kb scale a major technical challenge. Here, we present REWRITE, a modular platform for scar-minimized genome writing that enables engineering of large human genomic loci exceeding 100 kb at their native chromosomal sites. Using REWRITE, we deleted 105 kb spanning the dispersed HLA class-I locus and installed either compact or full-length refactored synthetic HLA haplotypes. Both refactored architectures exhibited transient activity before re-acquiring the native inactive state; upon extended culture, engineered lines yielded global transcriptomes nearly indistinguishable from parental cells. The engineered loci were genomically stable, retained interferon-responsive HLA expression, and remained compatible with differentiation into endothelial-like and thymic epithelial-like cells. Together, these results establish REWRITE as a platform for locus-scale genome engineering in hPSCs and provides a system for studying HLA haplotype structure and function in an isogenic pluripotent cell background.

**One sentence:** A practical platform for scar-minimized rewriting of native human loci exceeding 100 kb in hPSCs, demonstrated through synthetic refactoring of the class-I HLA region.

## Introduction

Human leukocyte antigen (HLA) class-I molecules govern antigen presentation and immune recognition across most cells ^1^. They are thus central to immune rejection, immune education, tolerance, and context-specific immune interactions ^2^. Moreover, the HLA locus is among the most consequential regions in the human genome, with strong associations to immune-related diseases and transplantation outcomes ^2–4^. It is also among the most polymorphic gene families in the genome ^5^, particularly the class-I genes *HLA-A, -B*, and *-C*, which are expressed by nearly all nucleated cells ^6^. This genetic diversity creates a compatibility barrier for allogeneic cells, as mismatched HLAs impair immune tolerance and limit applications requiring precise immunological interaction.

In many biological and engineering contexts, however, immune function depends on accurate specification of HLA haplotypes rather than elimination of HLA expression ^1^. Existing approaches to HLA incompatibility include HLA-homozygous haplotype banks ^7, 8^, which reduce mismatch but scale poorly, and “hypoimmune” strategies that suppress or delete HLA expression to evade immune rejection ^9–13^. While valuable for some applications, these approaches are not suited to settings in which HLA-mediated immune recognition must be preserved, such as macrophages for tissue surveillance, dendritic cells for antigen presentation, and thymic epithelial cells for T-cell education ^14–16^. Therefore, the present work focuses on enabling programmable specification of HLA haplotypes in a suitable “chassis” cell line.

Despite this need, targeted reconstruction of native HLA loci remains technically challenging. HLA regions span tens to hundreds of kilobases and comprise tightly coupled regulatory and coding architecture ^6^, placing them beyond the practical limits of conventional genome editing approaches. The 3.6-megabase HLA super-locus evolved through rapid gene-family expansion ^17^, resulting in high sequence similarity and repeat density ^18^ that complicate precise manipulation using homology-directed repair–based technologies. Together, these features make the class-I HLA locus an especially stringent test case for native-locus genome engineering in human pluripotent stem cells (hPSCs).

Recently, “genome writing” technologies have enabled construction of fully synthetic chromosomes in single-celled organisms, integrating DNA synthesis, in vitro assembly, and in vivo editing ^19^. In mammalian systems, “bigDNA” integration strategies in mouse embryonic stem cells (mESCs) have installed synthetic loci >100 kb in size while preserving native regulatory elements for developmental or disease modeling ^20–27^. However, translating these approaches to hPSCs remains a major challenge. Compared to mESCs, hPSCs are both less tractable and more fragile: they have lower transfection efficiency, reduced homologous recombination, increased p53-mediated apoptosis, and anoikis when cultured as single cells ^28–31^. Site-specific integration of smaller constructs (~10 kb) has been reported in hPSCs ^32^, and one report demonstrated a 173 kb bacterial artificial chromosome (BAC) ectopic integration ^33^. However, scar-minimized rewriting of native human loci exceeding 100 kb in hPSCs has remained a major barrier.

Here, we demonstrate scar-minimized refactoring of a native human genomic locus at >100 kb scale in hPSCs. To do so, we developed **REWRITE** (**RE**combinase **W**riting by **R**epeatable **I**ntegrations and **T**ag **E**xcision), a system for native locus-scale engineering in hPSCs. We applied REWRITE to reconfigure the human class-I HLA locus by installing synthetic haplotypes in either compact or full-length architectures. This establishes a practical benchmark for locus-scale genome engineering in hPSCs and provides a framework for systematic interrogation of complex allele-specific loci within their native genomic context.

## Results

### REWRITE enables genome rewriting in hPSCs

Locus-scale genome writing in hPSCs is constrained by cumulative DNA damage, transcriptional silencing, payload instability, and cell death ^28–30^, which together limit payload size and durability in existing landing pad (LP) architectures. To enable reproducible construction and integration of >100 kb synthetic payloads in hPSCs, we developed REWRITE, a staged workflow in which a recombinase-compatible landing pad is first installed and subsequently used for site-specific payload exchange and marker excision ^29^.

REWRITE is built around a “promoter dock” landing pad architecture that activates incoming payloads only after precise recombinase-mediated cassette exchange (RMCE), thereby minimizing background integration (Fig. 1a). The landing pad contains a fixed EF1α promoter, start codon, and loxP site that functions as a reusable “promoter dock”. The payload vector lacks a promoter and start codon but carries alternative selection markers that become transcriptionally active only upon in-frame RMCE. The payload vector also encodes an inducible Flp-ERT2 module (“FlpOut”) and a second FRT site, enabling tamoxifen-inducible excision of selection markers and generation of scar-minimized final payloads.

**Fig. 1.**
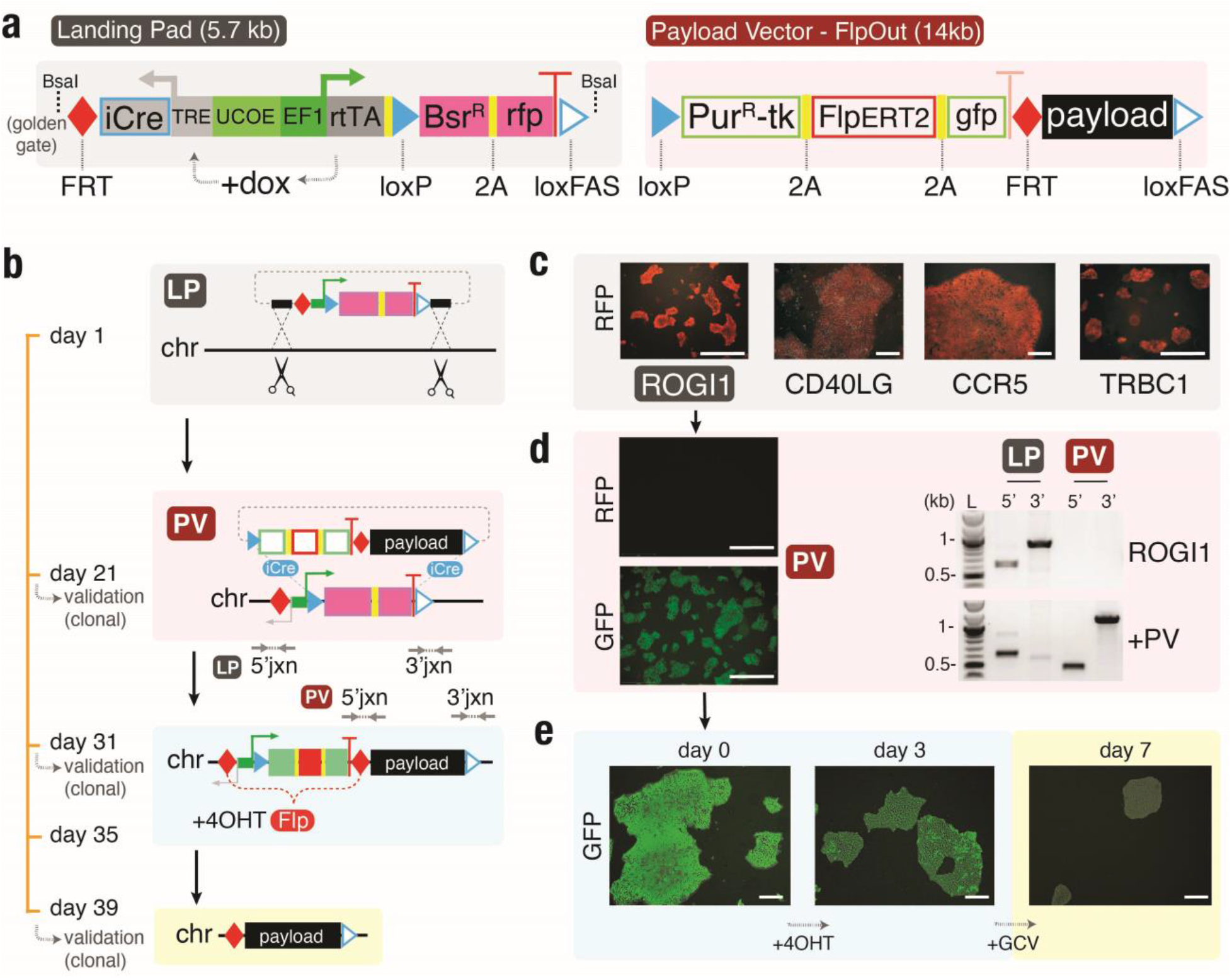
REWRITE platform architecture in hPSCs. (**a**) Schematic of the Landing Pad (LPneo) and terminal Payload Vector (FlpOut/PV). LPneo contains a doxycycline-inducible codon-optimized iCre (via TRE-rtTA), Blasticidin resistance (Bsr^R^), mScarlet (RFP), and a wild-type loxP site, flanked by an FRT site for marker removal. An alternate version (LPcore) lacks TRE-iCre. Golden Gate cloning is used to append locus-specific homology arms via BsaI cut sites. FlpOut vectors encode a puromycin–thymidine kinase fusion (Pur^R^-TK), tamoxifen-inducible FlpERT2, mNeonGreen (GFP), and a second FRT site. (**b**) Timeline schematic illustrating LPneo insertion, Payload integration, and FlpOut inducible marker removal. Each step is a single-cell derived clone, with the rate-limiting step being cell expansion time. (**c**) Targeted integration of LPneo into four transcriptionally silent loci in hPSCs, with junction PCR confirming successful LPneo insertion. (**d**) RMCE-mediated integration of FlpOut into the LPneo-ROGI1 locus, showing RFP-to-GFP marker conversion and junction PCR validation. (**e**) Tamoxifen induction of FlpERT2 results in >90% marker excision by day 3, with additional negative-selection using ganciclovir (GCV) to remove residual Pur^R^-TK–positive cells.

This configuration ensures that off-target integrations are largely inert, while correctly recombined payloads become transcriptionally active, sharply reducing background compared to LP systems in which payloads carry their own active promoters.

We benchmarked prototype LP designs at the ROSA26 locus in mouse embryonic stem cells (mESCs) (Supplementary Fig. S1a–b). We first assessed the importance of Cre availability, confirming that co-nucleofection of a codon-optimized iCre (which improves translation in human cells) increased colony yield for integration of a 10 kb *HLA-A* payload eightfold compared to wild-type Cre (Supplementary Fig. S1c–f). Consistent with a prior report, the heterotypic loxP/loxFAS pairing outperformed loxP/lox2272 by ~15-fold in RMCE efficiency ^34^. However, progressive LP silencing in the absence of blasticidin selection led us to incorporate a Universal Chromatin Opening Element (UCOE) ^35^, which maintained mScarlet expression for >6 days without selection (Supplementary Fig. S2).

These optimizations yielded an LP design (“LPneo”) that incorporates iCre under a doxycycline-inducible TRE3G promoter, the reverse tetracycline-controlled Transcriptional Activator (rtTA) gene, relocated loxP positioning for in-frame docking, and Golden Gate sites for modular homology arm assembly, enabling deployment to additional genomic loci (Fig. 1a). A simplified variant (“LPcore”), lacks the inducible iCre, instead utilizing transient plasmid-expressed iCre.

To assess performance at different genomic sites, we integrated LPneo at four transcriptionally inactive (in hPSCs) loci in PGP1 hPSCs (CD40LG, TRBC1, ROGI1, and CCR5), the latter two being common safe-harbor sites (Fig. 1c, Supplementary Fig. S3a), using CRISPR-assisted HDR. Junction PCR, alkaline phosphatase staining, and RT-qPCR confirmed correct integration, maintenance of pluripotency, and minimal LPneo silencing over 29 days extended culture (Supplementary Fig. S3b-e). Single-cell derived clonal LP lines expanded to usable cell densities within ~21 days.

We next tested integration of the “FlpOut” payload vector (PV) in ROGI1-LPneo hPSCs (Fig. 1d). We found that co-nucleofecting anti-apoptosis plasmid *BCL-XL* increased yields from 24 to 104 colonies per million cells. Junction PCR confirmed complete RMCE and loss of the original LP cassette for four selected clones (Fig. 1d, Supplementary Fig. S4a). Clonal integrants reached densities suitable for marker excision within 7–10 days.

Following tamoxifen induction to evaluate marker removal, excision efficiencies ranged from 50– 90% within 3 days, depending on colony size (Fig. 1e, Supplementary Fig. S4b–c). Negative selection against residual markers further enriched excised clones. PCR and Sanger sequencing confirmed marker removal, (Supplemental Fig. S4d–e). Thus, FlpOut enables marker excision within the same workflow without secondary transfection or clonal outgrowth.

Across these stages, LP installation, payload integration, and marker excision — each requiring clonal expansion — could be completed within ~39 days when DNA constructs are available, with each stage yielding a stable, clonal intermediate. The primary rate-limiting step of the workflow timeline is clonal cell expansion at each stage to transfectable densities necessary for the subsequent stage. Together, these results show that promoter docking, optimized lox pairing, anti-silencing elements, inducible cassette excision, and transient survival support collectively enable reproducible locus engineering in hPSCs.

### synHLA: a refactored class-I locus for designer HLA haplotypes

Because the class-I HLA locus combines megabase-scale dispersion, extreme polymorphism, and dense regulatory sequences, successful refactoring here represents a stringent test case for locus-scale engineering in hPSCs. To enable single-step installation of full class-I HLA haplotypes, we engineered a synthetic, refactored locus (“synHLA”) consolidating targeted sequences into a contiguous insert, thereby avoiding their dispersal across a megabase ^36^. In the native genome, *HLA-B* and *-C* are separated by 80 kb, and *HLA-A* lies 1.3 megabases upstream (Fig. 2a). We captured the 93 kb *HLA-B/C* region from a linearized BAC into the FlpOut-PV backbone by yeast recombineering, then inserted *HLA-A* with all its ENCODE-predicted enhancers (Supplementary Fig. S5a; PCR-amplified in two 5 kb fragments) 36 kb downstream of *HLA-B* (Fig. 2b–c), a region lacking annotated enhancers or accessible chromatin in available ENCODE datasets. Deep sequencing verified the assembly (Supplementary Fig. S5b). The resulting synHLA, a fully assembled 100 kb construct (115 kb with vector), encoded the rare haplotype A*30:01, B*55:01, and C*03:03.

**Fig. 2.**
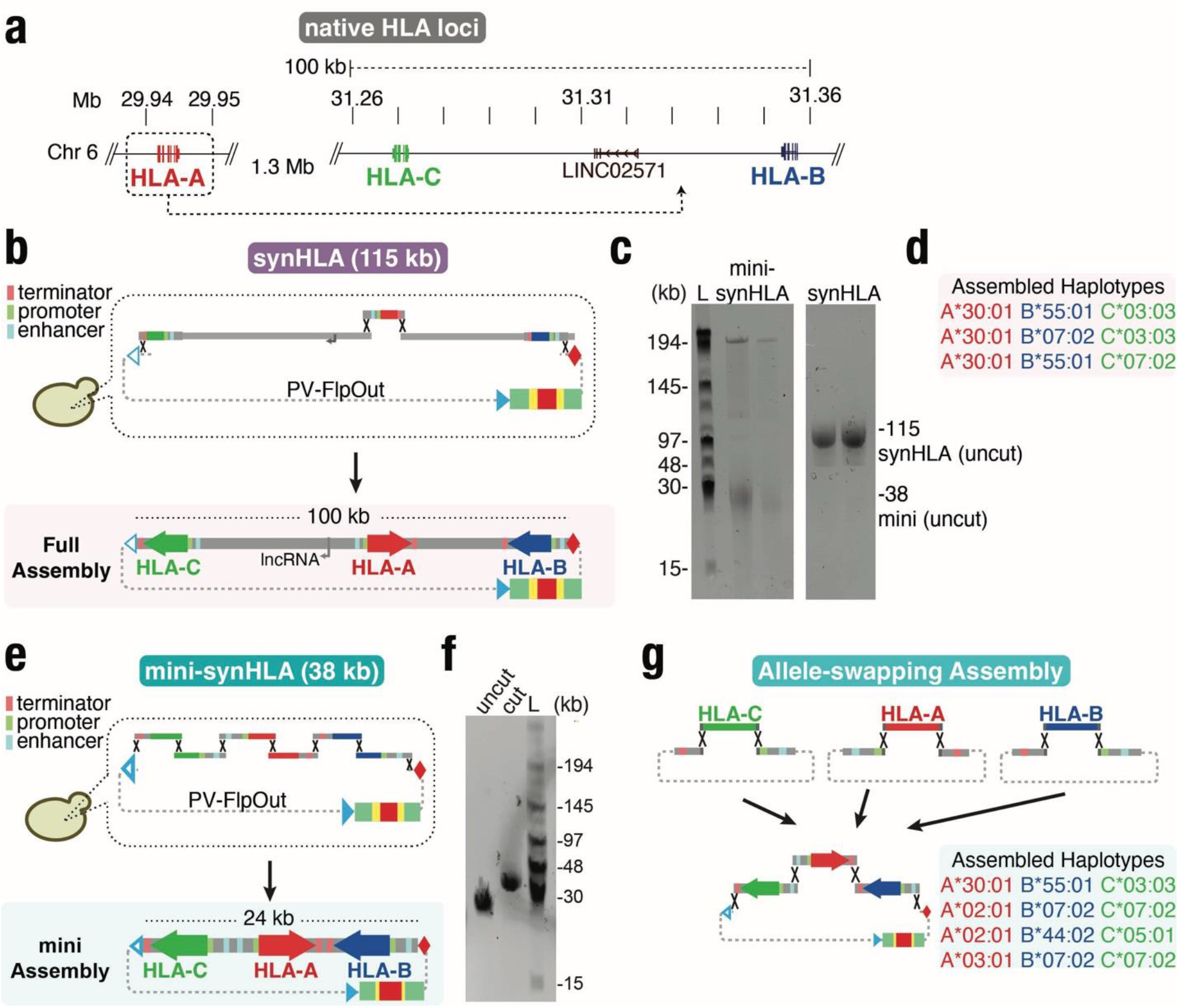
Synthetic HLA haplotypes assembled by modular bigDNA engineering. (**a**) Genomic organization of the native class-I HLA loci on chromosome 6, showing the 1.3 Mb separation between *HLA-A* and *HLA-B*. REWRITE enables repositioning of *HLA-A* into a contiguous synthetic haplotype with *HLA-B* and *HLA-C*. (**b**) Modular assembly of full-length synHLA (115 kb) by in-yeast recombineering using BAC-derived segments and synthetic junctions. FlpOut PV vector backbone shown. (**c**) Field-inversion gel electrophoresis (FIGE) of uncut synHLA and mini-synHLA constructs, confirming correct assembly and plasmid size integrity. (**d**) Example haplotypes generated by allele-swapping for class-I *HLA-A, -B*, and *-C* genes. Colors correspond to native loci shown in (A). (**e**)Modular assembly of compact mini-synHLA (38 kb) using defined enhancer, promoter, and coding segments for each gene. (**f**) FIGE validation of mini-synHLA construct size (uncut and restriction-digested forms). (**g**) Schematic of allele-swapping strategy using modular fragments to generate multiple synthetic haplotypes across different genetic backgrounds.

To test whether intergenic genomic context is required for function, we designed a compact variant (“mini-synHLA”) retaining only core regulatory loci. Enhancer boundaries were defined using ENCODE annotations, and intergenic DNA regions lacking annotated enhancers were excluded from the compact variant (Fig. 2e, Supplementary Fig. S5a). PCR-amplified genes (3–5 kb) were fused with native-sequence linkers and recombineered into the terminal PV, generating a 24 kb core insert (38 kb total) (Fig. 2e–f).

To customize haplotypes, we developed distinct “allele-swapping” strategies for synHLA and mini-synHLA (Fig. 2g, Supplementary Fig. S5d). For synHLA, whose size and repeat density precluded standard cloning ^37^, we used in-yeast recombineering with *URA3* counter-selection and CRISPR/Cas9-editing, yielding two additional rare haplotypes (Fig. 2d, Supplementary Fig. S5d). For mini-synHLA, we used a modular system of three vectors encoding full-length HLA loci, each edited and assembled using linker fragments via yeast recombineering. Final assemblies were validated by nanopore sequencing, generating three rare haplotypes (Fig. 2g, Supplementary Fig. S5c). These strategies support programmable construction of class-I HLA haplotypes in full-length or compact formats, with modular control over regulatory architecture and allele composition.

### Engineering biallelic class-I HLA deletions in hPSCs

To enable repeatable reconstitution of on-demand HLA haplotypes, we generated a panel of “blank” hPSC lines with biallelic deletions of the class-I HLAs, a technically demanding step due to the size and extensive polymorphism of the HLA region (Fig. 3a).

**Fig. 3.**
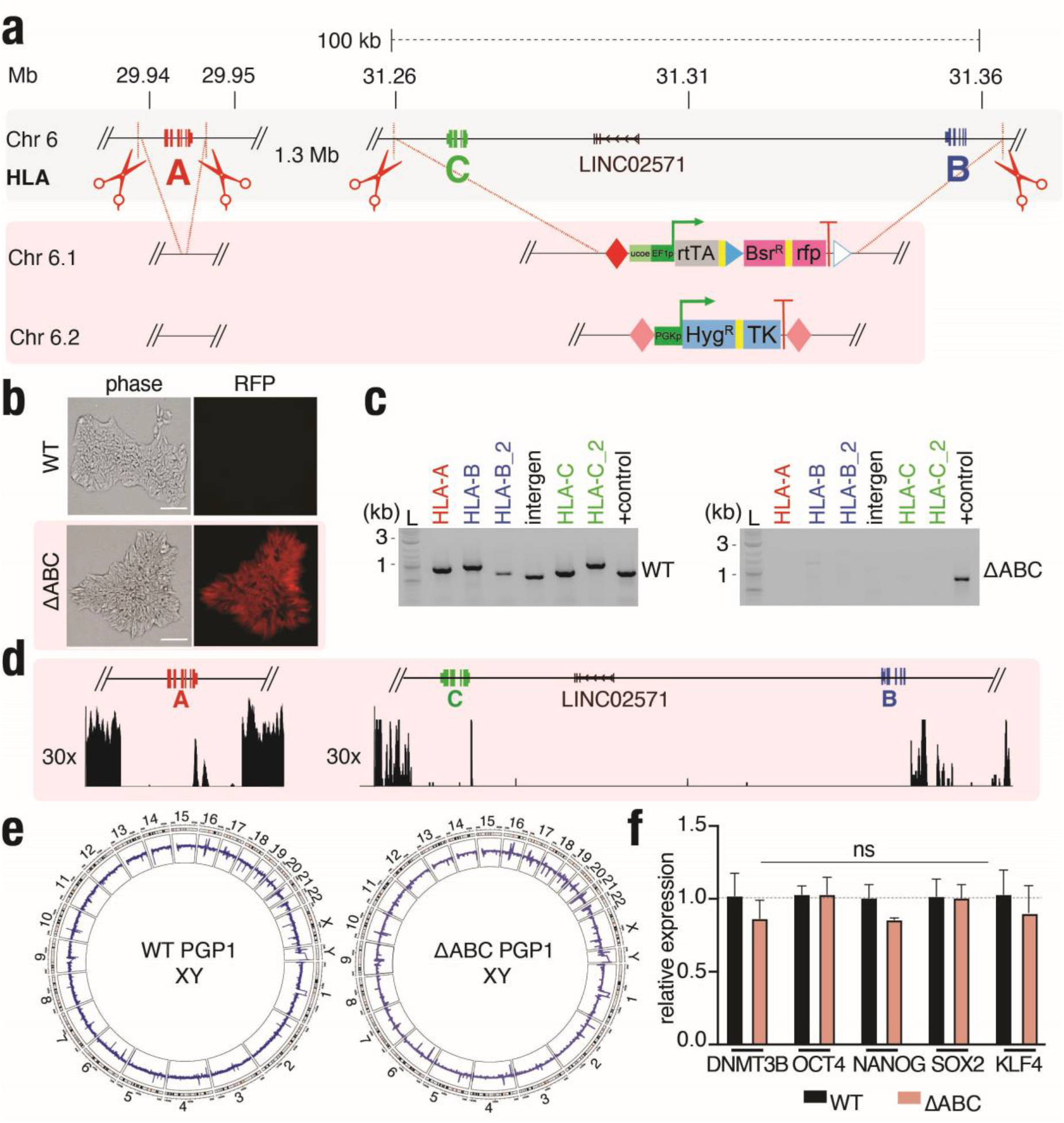
Bi-allelic removal of class-I HLA regions in human pluripotent stem cells. (**a**) CRISPR-based biallelic deletion of *HLA-A* (8 kb) and *HLA-B*/*C* locus (100 kb) in PGP1 hPSCs followed by landing pad (LPcore) and “Deleter” (HygTK) cassette integration. (**b**) Representative images of WT PGP1 and ΔABC hPSCs in phase contrast and with the landing pad integration (RFP) (Scale bar = 200 μm) (**c**) PCR gel verifying *HLA-A, -B*, -*C* class-I biallelic deletions in ΔABC. WT PGP1 as a control. (**d**) Coverage plots (hg38) at 30x depth whole-genome sequencing (WGS) showing biallelic deletion of both the HLA-A and HLA-B/C loci in PGP1 hPSCs. (**e**) Circos plot showing no chromosomal aberrations of PGP1-ΔABC. WT PGP1 as a control. (**f**) Expression of pluripotency genes was measured by RT-qPCR in WT PGP1 and class-I HLA ΔABC. Gene expression was normalized to WT PGP1 (dotted line). Data represent mean ± SEM from n=3 independent experiments and *p*-values were calculated by two-way ANOVA *t*-test (* *p*<0.05, ns=not significant).

We first deleted the ~8 kb *HLA-A* locus using a Cpf1-homolog, Mad7 (Supplementary Fig. S6a– d), using a scarless deletion-optimized genome editing strategy ^38^, which utilizes short single-stranded oligodeoxynucleotide (ssODN) templates as homology-directed repair (HDR) donors to specify the deletion, along with transient enrichment for transfected cells. Screening four gRNA pairs with 100 bp ssODN HDR donors, transient puromycin enrichment, and co-expressing the anti-apoptosis *BCL-XL* protein (which improves bi-allelic editing ^39^), yielded two sets with clean monoallelic deletions. Using the optimized set, we recovered a scarless biallelic deletion clone (PGP1-ΔA) from 168 randomly selected colonies and validated by whole genome sequencing (WGS) (Supplementary Fig. S6d).

After establishing the *HLA-A* deletion, we targeted the 93 kb *HLA-B/C* region, which was challenging due to repeat content and low gRNA specificity. We reasoned that the ability to utilize positive selection for the second deletion product would enable generation of the biallelic deletion. We constructed a “Deleter” construct expressing Hygromycin for positive selection and Thymidine Kinase for negative selection, flanked by mutant FRT sites for subsequent marker removal, and Golden Gate homology-arm assembly for deployment to specific sites. To improve biallelic deletion efficiency, we co-delivered Cas9 and Mad7 gRNAs with the Deleter construct (Supplementary Fig. S6e), LPcore for allelic integrations, anti-apoptosis expressing plasmid *BCL-XL* ^39^, and the non-homologous end-joining (NHEJ) inhibitor Nedisertib to bias toward homology-directed repair. Nucleofection of a million PGP1 or PGP1-ΔA cells yielded 30–50 colonies, the majority with at least monoallelic deletions. Across multiple experiments, we obtained several clones achieving full biallelic deletion (PGP1-ΔBC), and multiple (four) independent fully “blank”-HLA lines (Fig. 3b–c, Supplementary Figs. S6f–g, S7a). Downstream studies focused on one of those clones (PGP1-ΔABC).

We validated genomic integrity via 30x depth WGS in four distinct blank-HLA clones, with a particular focus PGP1-ΔABC. All showed the expected deletions, no structural abnormalities or recurrent pathogenic *TP53* mutations ^40^ (Fig. 3d, Supplementary Figs. S6h, S7b). Euploidy in PGP1-ΔABC was independently confirmed by WGS (Fig. 3e). To evaluate the impact of large biallelic deletions on pluripotency, we performed RT-qPCR on blank-clone PGP1-ΔABC, which had tightly packed typical primed hPSC morphology (Fig. 3b). This clone retained high expression of pluripotency markers (Fig. 3f), confirming that the genetic manipulations did not alter pluripotency potential.

Thus, we established a validated class-I HLA-blank clonal hPSC line suitable for reconstitution of synthetic HLA haplotypes.

### REWRITE enables integration of >100 kb synthetic loci at native sites

To test whether synthetic, refactored HLA loci could be installed and function within their native genomic context, we delivered both mini-synHLA (38 kb) and full-length synHLA (115 kb) constructs into the validated fully class-I–deleted hPSC line (PGP1-ΔABC). Both constructs were introduced into the REWRITE landing pad by iCre-mediated recombination in PGP1-ΔABC (Figs. 1, 4b, Supplementary Fig. S8a–b).

**Fig. 4.**
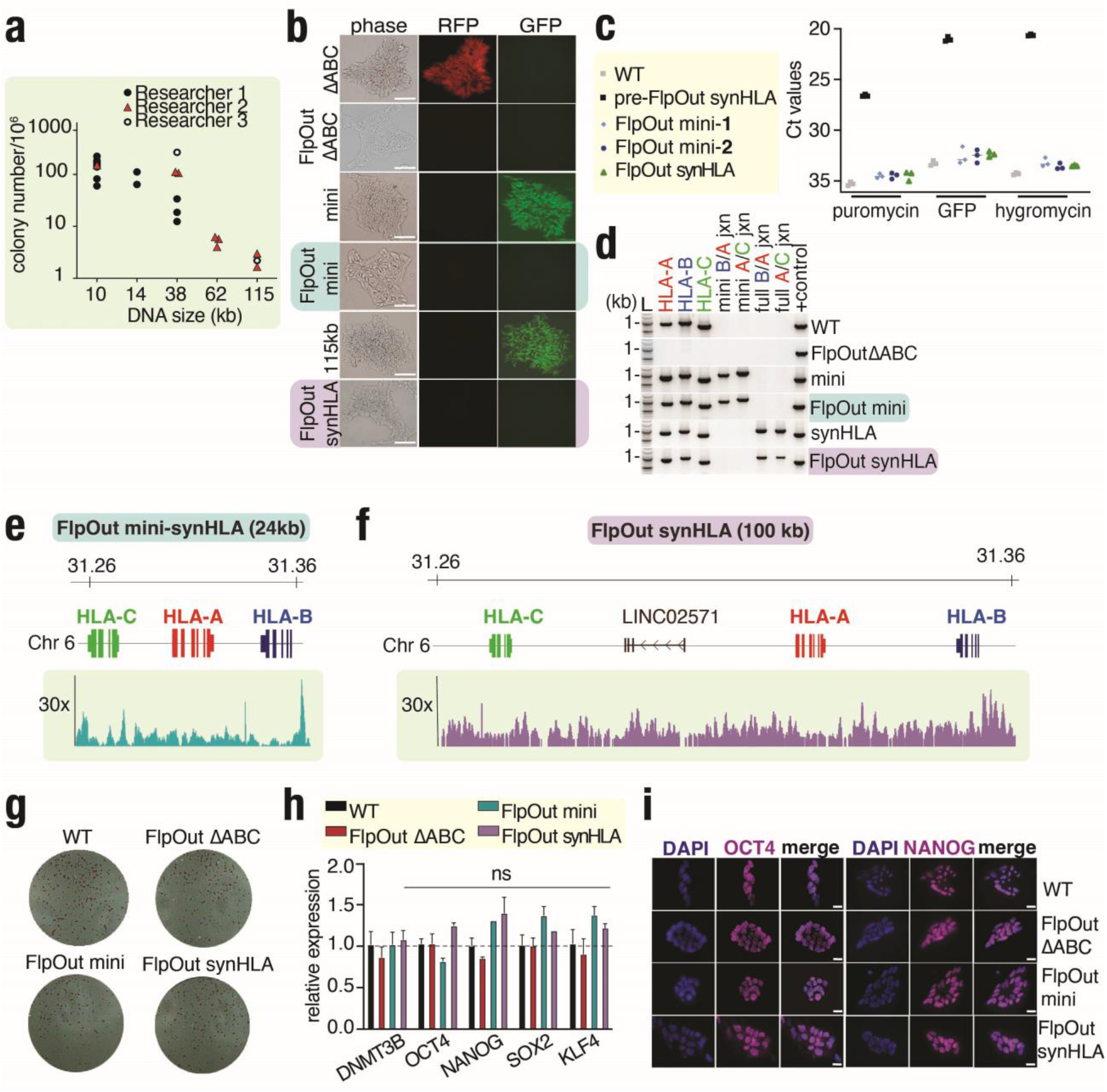
Genome rewriting of class-I HLA region in human pluripotent stem cells. (**a**) Total integrated colonies obtained normalized per million cells nucleofected for different DNA size (kb) payloads using the REWRITE method. The graph shows experiments performed by three different researchers. (**b**) Representative images of ΔABC, mini- and synHLA integration before and after flippase removal of the cassette are shown in phase contrast, with the landing pad integration (RFP), and the payload deliveries (GFP). (Scale bar = 200 μm). (**c**) Verification of complete marker excision in all FlpOut hPSCs by qPCR. Pre-excision synHLA hPSCs with a Ct value in the 20s. All other samples showed non-specific detection. Data represent mean ± SEM from n = 3 independent experiments. (**c**) PCR gel verifying the integration of the class-I HLA genes upon delivery of mini- and synHLA payloads. WT PGP1 and ΔABC as controls. Specific junctions (jxn) between HLA-B/A and HLA-A/C are amplified to show the integration of the synthetic cassette. (**e-f**) The precise delivery of class-I HLA genes for both mini- and synHLA payloads into the native site is shown via WGS against hg38. (**g**) Images representing alkaline phosphatase staining for each marker-excised (FlpOut) cell line. WT PGP1 as a control. (**h**) Expression of pluripotency genes was measured by RT-qPCR in WT PGP1, FlpOut ΔABC, FlpOut mini-synHLA and synHLA. Gene expression was normalized to WT PGP1 as a control (dotted line). Data represent mean ± SEM from n=3 independent experiments. *P*-values were calculated by two-way ANOVA *t*-test (* *p*<0.05, ns = not significant). (**i**) Immunostaining images of OCT4 and NANOG in WT PGP1, FlpOut ΔABC, FlpOut mini- and synHLA. Cell nuclei were stained with DAPI. (Scale bar = 50 μm).

Across multiple independent experiments and users, we consistently observed a strong size-dependent practical integration limit. Mini-synHLA (38 kb) yielded approximately 100 selected colonies per million nucleofected cells, whereas full-length synHLA (115 kb) yielded approximately 1.5–3 colonies per million cells. However, experiments were routinely scaled (up to 5 million cells), enabling recovery of multiple correctly integrated clones (Fig. 4a). We also tested an intermediate-size (62 kb) synthetic payload assembled from a nearby human genomic region to further define the size-dependence of integration efficiency, which yielded intermediate colony numbers (Fig. 4a). This size-dependent drop-off is consistent with known biophysical constraints on large DNA delivery in hPSCs ^41^. Notably, the observed limit is consistent with the only comparable prior report, in which a 173 kb BAC integration at an ectopic site yielded a single recovered clone ^33^. The approximate relationship between payload size and recovered integration events (Fig. 4a) therefore provides a practical benchmark for bigDNA integration in hPSCs. Nevertheless, REWRITE reproducibly yielded clonally isolated, fully validated integrants in every experiment, with all recovered clones exhibiting precise, full-length integration at the intended genomic site and no detectable background or partial insertions.

Although integration frequency decreased with payload size, correctly integrated clonal lines were reproducibly recovered in every experiment. In these clones, the marker-cassette was completely excised on both alleles using the FlpOut system, as observed by loss of fluorescence (Fig. 4b). We confirmed complete excision of the UCOE/EF1α cassette using the FlpOut system by RT-qPCR and 30x WGS (Figs. 4c, Supplementary Fig. S9a), confirming that only refactored native sequences remained. Next, we validated the genomic integrity of our clones, performing tiled internal PCR and WGS to confirm full-length integration, no off-target integrations or pathogenic *TP53* mutations, and absence of off-target structural variation, gross genomic abnormalities, or aneuploidies (Fig. 4d, 4e-f, Supplementary Fig. S9b). Engineered clones retained high pluripotency across early and late passages (Fig. 4g, 4h, 4i, Supplementary Fig. S8c–d),

To assess the generality of bigDNA delivery, we also placed LPcore at a second transcriptionally silenced location on chromosome 6 approximately 1.1 Mb from the class-I HLAs in a female hPSC line (NCRM2) (Supplementary Fig. S10a-d). Deliveries of both mini-synHLA and full synHLA to this hPSC background were at similar frequencies as for that into the HLA-blank male hPSC PGP1 (Supplementary Fig. S10e, S10f), and this maintained high pluripotency (Supplementary Fig. S10g, S10h).

Together, these results demonstrate that REWRITE enables precise installation of >100 kb synthetic loci at native genomic sites in hPSCs while maintaining genome stability and pluripotency consistent with established International Society for Stem Cell Research (ISSCR) standards ^40, 42, 43^. A staged workflow, low-background promoter docking, and inducible marker excision collectively supported recovery of full-length, correctly integrated payloads even at large DNA sizes, overcoming key practical limitations of existing landing pad systems.

### Synthetic HLA loci are transiently overexpressed before resolving to a wild-type state

Installation of large synthetic loci at native genomic sites introduces “naked DNA” and regulatory elements that have not participated in the developmental history of the host genome. How such refactored loci behave transcriptionally following integration — particularly in a pluripotent context where the native HLA region is normally inactive or minimally expressed ^44^ — is thus an open question. To examine activation status following locus-scale rewriting, we assessed basal HLA expression at the single-cell level in clonally derived marker-excised (FlpOut) hPSCs by flow cytometry using the EMR8-5 antibody — which detects total cell-surface HLA heavy chain protein density independently of its conformational or complex-assembled state (Fig. 5a) ^44^.

**Fig. 5.**
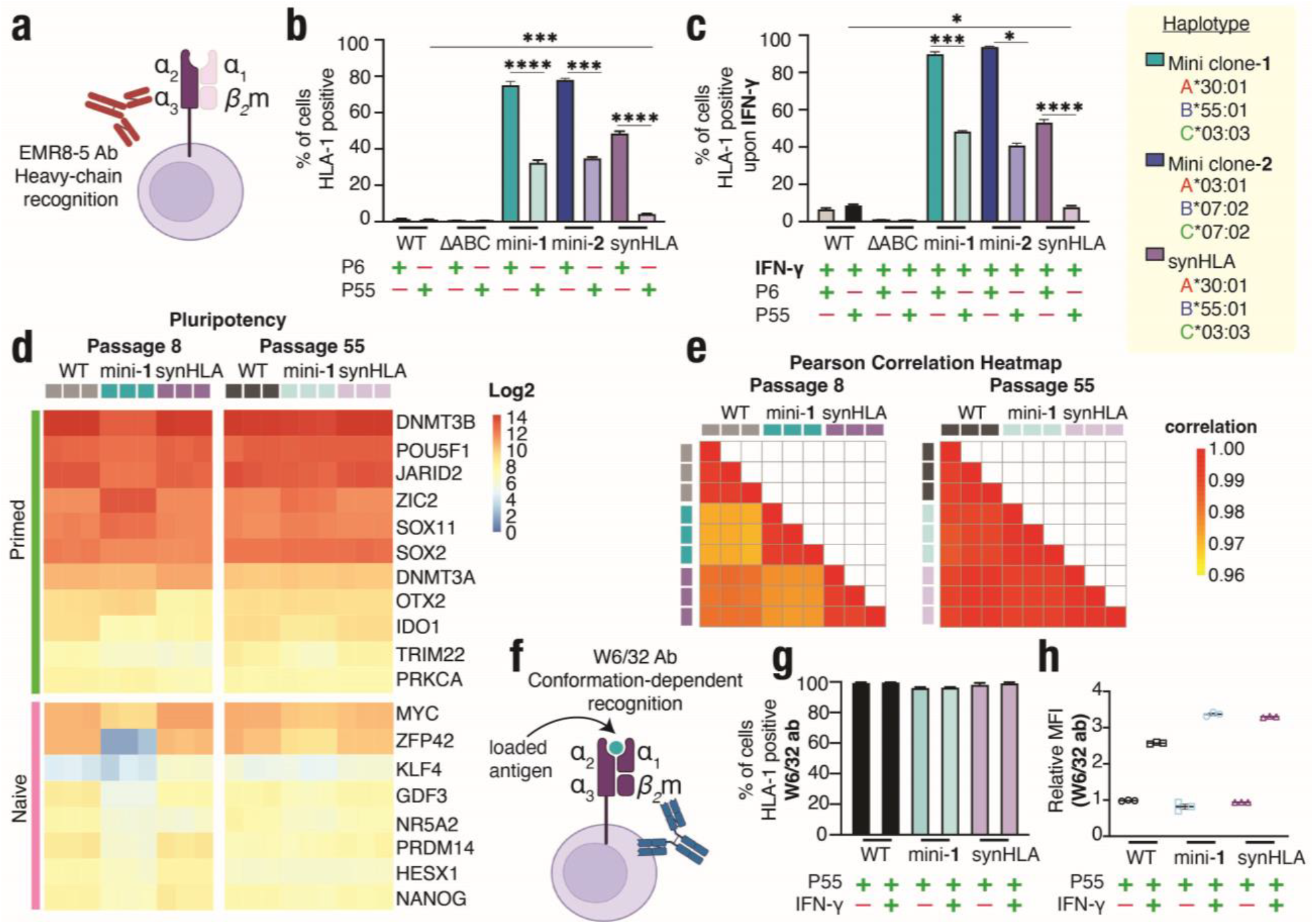
Restoration of HLA expression reveals cell state constraints and resolution. (**a**) Schematic representation of HLA class-I recognition by EMR8-5 antibody. EMR8-5 detects the total amount of HLA class-I heavy chains regardless of native complex conformation. (**b**) The expression of class-I HLA was determined by flow cytometry in WT PGP1, FlpOut ΔABC, FlpOut (marker-excised) mini-1 and mini-2, and synHLA hPSCs without interferon-<u>γ (I</u>FN-γ) induction at early passage (P6, passage 6) and after 55 additional passages (P55, passage 55). Class-I HLA is stained with anti-HLA class-I antibody (EMR8-5 clone). Data represent mean ± SEM from n = 3 independent experiments. *P*-values were calculated by unpaired two-tailed *t*-test (\**p*<0.05, ** < 0.01, ***< 0.001, ****<0.0001, ns= not significant). Two different haplotypes for the FlpOut mini-synHLAs were used (shown in the table). (**c**) The expression of class-I HLA was determined by flow cytometry in WT PGP1, FlpOut ΔABC, FlpOut (marker-excised) mini-1 and mini-2 and synHLA hPSCs upon interferon-<u>γ (I</u>FN-γ) induction at early passage (P6, passage 6) and after 55 additional passages (P55, passage 55). Class-I HLA is stained with anti-HLA class-I antibody (EMR8-5 clone). Data represent mean ± SEM from n = 3 independent experiments. *P*-values were calculated by unpaired two-tailed *t*-test (\**p*<0.05, ** < 0.01, ***< 0.001, ****<0.0001, ns= not significant). Two different haplotypes for the FlpOut mini-synHLAs were used (shown in the table). (**d**) Log_2_ gene expression heatmap of pluripotency markers from RNA-seq analysis of WT, FlpOut (marker-excised) mini-1, and synHLA hPSCs at passage 8 and passage 55. (**e**) Pearson correlation analysis of WT-, FlpOut (marker-excised) mini-1-, and synHLA hPSC at passages 8 and passage 55. Pairwise Pearson correlation coefficients (r) are displayed as heat maps. All samples demonstrated strong correlations (r = 0.96–1.00) across passages. (**f**) Schematic representation of HLA class-I recognition by W6/32 antibody. W6/32 detects surface-folded HLA class-I heavy chain/β2-microglobulin/loaded antigen complexes. (**g**) Cell surface HLA class-I expression was assessed by flow cytometry using W6/32, a conformation-dependent monoclonal antibody specific for properly assembled HLA class-I heavy chain–β2-microglobulin complexes. The expression was assessed at Passage 55 (P55) in WT PGP1, FlpOut ΔABC, FlpOut (marker-excised) mini- and synHLA hPSCs with and without IFN-γ induction. Data represent mean ± SEM from n = 3 independent experiments. (**h**) The graph shows the median of fluorescence intensity (MFI) normalized to the WT PGP1 control determined by flow cytometry in WT PGP1, FlpOut mini- and synHLA hPSCs with and without IFN-γ induction at passage 55 (P55). Background median fluorescence intensity from the secondary-only control was subtracted from all samples (ΔMFI). Data are expressed as fold change in ΔMFI over the WT PGP1 control. Class-I HLA was stained with anti-pan HLA class-I antibody (W6/32 clone). Data represent mean ± SEM from n = 3 independent experiments.

As expected, wild-type hPSCs exhibited negligible class-I HLA expression (<1%) (Fig. 5b) ^44^. In contrast to wild-type, both mini- and full-length synHLA loci revealed significant expression, with approximately 78% and 50%, respectively, of cells expressing class-I HLA at passage 6. The mini-synHLA showed higher baseline expression and long-term stability, maintaining 45% expression at passage 55 (Fig. 5b). Notably, two independent mini-synHLA clones with distinct haplotypes exhibited similar HLA expression over time.

By contrast, the full length synHLA showed greater progressive silencing, dropping to just 5% by passage 55, resembling the baseline expression level of unstimulated parental wildtype hPSC (Fig. 5b). A multi-passage time course (P6, P20, P35, P45, and P55) showed a passage-dependent decrease in the percentage of cells expressing HLA, suggesting a gradual convergence of the synthetic expression levels towards the native low-expression baseline (Supplementary Fig. S11a). A linear regression analysis confirmed the passage-dependent decline in class-I HLA expression in both cell lines exhibiting a similar R^2^ coefficient, indicating a predictable rate of loss over time (Supplementary Fig. S11c).

Importantly, flow cytometry analysis confirmed that cell viability is uniform and high across all engineered cell lines under both basal and IFN-γ stimulated conditions, confirming that the differential HLA expression was not due to preferential cell survival or clonal selection (Supplementary Fig. S11d). Both synthetic constructs remained highly responsive to IFN-γ across all the passages (Fig. 5c, Supplementary Fig. S11b), reinforced by RNA-seq transcriptomic heatmaps confirming the activation of interferon response genes (Supplementary Fig. S12b). Finally, RNA-seq profiling showed that the robust expression of the mini-synHLA is not caused by improper transcription or activation of the neighboring genes (+/– 200 kb) surrounding the engineered HLA locus (Supplementary Fig. S11e).

To further confirm that neither the genome writing nor the extended passaging were compromising stem cell identity, we analyzed the global transcriptomic profile via RNA-seq ^45^. Log-normalized expression heatmaps revealed no significant differences in the expression of pluripotency genes, for both naive and primed state markers, at both early and late passages across different cell lines (Fig. 5d). Moreover, expression levels of genes representing the three germ layers were comparably low to those of parental PGP1 hPSCs (Supplementary Fig. S12a). Global transcriptomic stability was further confirmed by a Pearson correlation heatmap, comparing P8 and P55 states, which revealed a high correlation range of 0.96–1.00, converging on a transcriptional state nearly indistinguishable from wild-type PGP1 hPSC (Fig. 5e). This convergence on the parental baseline suggests that the REWRITE platform enables locus-scale engineering without inducing permanent shifts in cell identity or global gene expression.

Next, we evaluated the structural integrity of our synthetic class-I HLA proteins by staining the cells with the conformation dependent W6/32 antibody at P8 and P55. While EMR8-5 allowed us to quantify total synthetic HLA heavy chain accumulation, it does not assess structural fidelity. The W6/32 antibody strictly recognizes a fully assembled trimeric complex consisting of HLA heavy chain, beta-2-microglobulin (β_2_M), and an endogenously loaded antigenic peptide (Fig. 5f), thereby establishing whether the synthetic loci generate properly assembled surface complexes ^46–48^. Of note, W6/32 is extremely sensitive to even low levels of properly assembled HLA complex. As expected, WT hPSCs have a low-density yet uniform population of fully surface assembled class-I HLA complexes detected as nearly 100% positive by W6/32, despite having an absolute heavy-chain density too low to be recognized by the sequence-specific EMR8-5 antibody.

Using W6/32, we found that both mini- and full-synHLA produced stable peptide-loaded class-I HLA complexes that mirrored the WT hPSC condition, indicating successful processing through the native antigen-presentation pathway despite the passage-dependent decrease in absolute expression levels (Fig. 5g, Supplementary Fig. S11f) ^46–48^. Notably, the relative Median Fluorescence Intensity (MFI) confirmed that both cell lines at late passage recapitulated the functional responsiveness of the WT hPSC (Fig. 5h).

Together, these results show that locus-scale genome rewriting can transiently generate non-native activation states, which progressively resolve toward the native transcriptional baseline with extended culture without compromising stem cell identity. Moreover, these observations suggest that a compressed locus functions similarly to a full-length locus that contains additional native non-regulatory intergenic DNA. Since mini-synHLA haplotypes are simpler to generate and have higher integration frequencies, these results support the compact architecture as a practical alternative for generating isogenic HLA-specific hPSCs.

### Engineered hPSCs differentiate to defined lineages

Finally, we asked whether locus-scale genome rewriting preserved the ability of engineered hPSCs to differentiate into defined lineages using either transcription factor–based forward programming or conventional directed differentiation. In parallel, we sought to determine whether the synthetic loci remained responsive to lineage-specific regulatory programs.

We first differentiated FlpOut full- and mini-synHLA engineered hPSCs at late passage (P55) into mesodermal endothelial-like cells (EC) using more stringent conventional directed differentiation, with a streamlined 8-day protocol (Fig. 6a). Following differentiation, immunostaining confirmed expression of the endothelial marker CD31 showing no differences with the parental cell line (Fig. 6b). Upon lineage commitment, both the mini- and full-synHLA hPSC-ECs showed a robust gain of basal class-I HLA expression, largely mirroring the upregulation observed during the wildtype hPSC differentiation kinetics (Fig. 6c). Moreover, there was a highly significant increase in the relative MFI when comparing P55 undifferentiated hPSCs directly to their differentiated EC counterparts, suggesting a greater density of HLA (Fig. 6d).

**Fig. 6.**
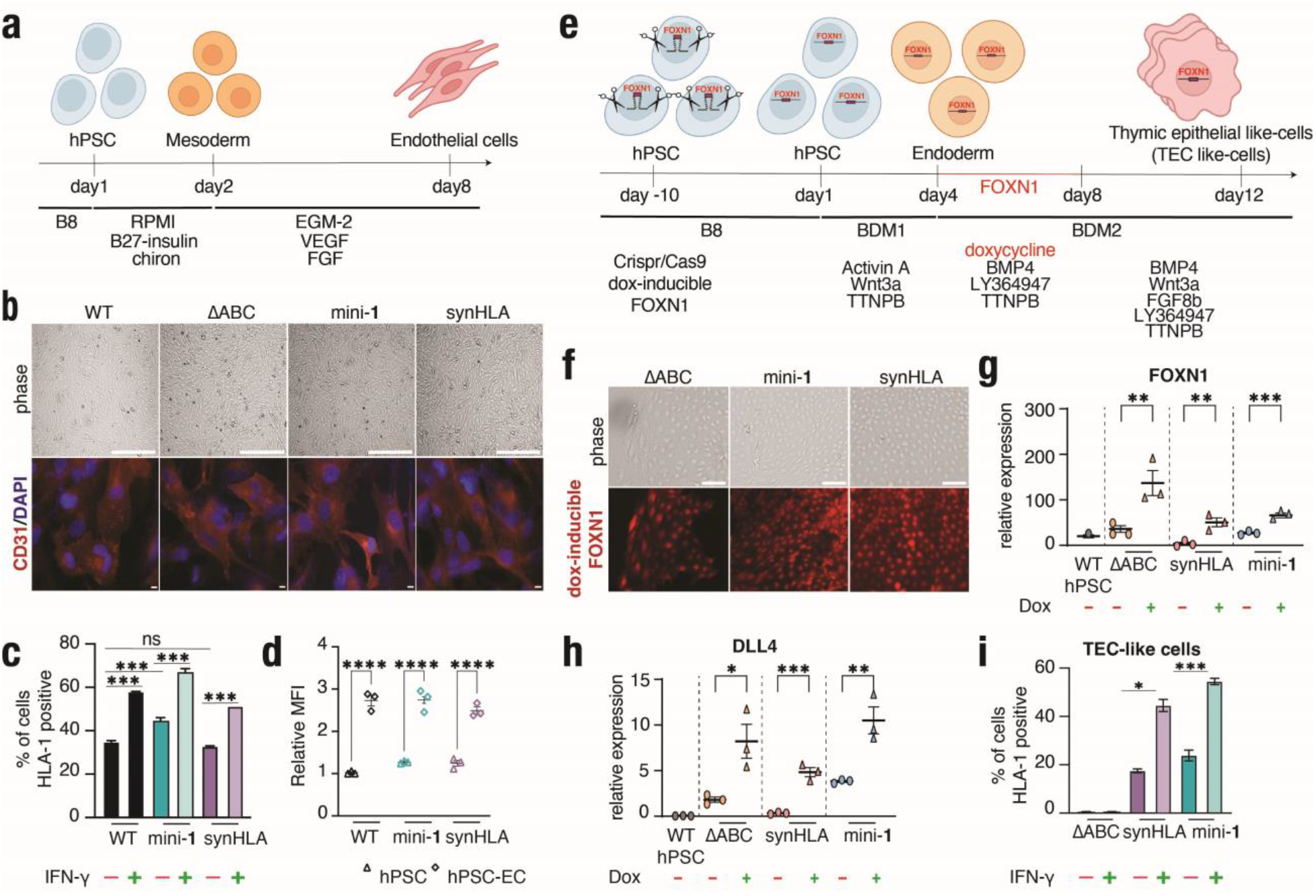
Differentiation of HLA engineered hPSCs. (**a**) Schematic diagram of the experimental protocol to differentiate hPSCs to endothelial cells (EC). (**b**) Representative images of WT PGP1, FlpOut ΔABC, FlpOut mini-1 and synHLA hPSC upon mesodermal endothelial cell differentiation are shown in phase contrast (upper panel) (scale bar = 200 μm). Lower panel: WT PGP1, FlpOut ΔABC, FlpOut mini-1 and synHLA hPSC were stained with EC marker CD31 marker (red) and DAPI. (scale bar=50 μm). (**c**) The expression of class-I HLA was determined by flow cytometry in WT PGP1, FlpOut ΔABC, FlpOut mini-1 and synHLA hPSCs-EC derived with and without interferon-<u>γ (I</u>FN-γ) induction at passage 55. Data represents mean ± SEM from n = 3 independent experiments. *P*-values were calculated by one-way ANOVA *t*-test (*** *p*<0.001, ns = not significant). (**d**) The graph shows the median of fluorescence intensity (MFI) normalized to the WT PGP1 control determined by flow cytometry in WT PGP1, FlpOut mini- and synHLA hPSCs and hPSC-ECs derived at passage 55 (P55). Background median fluorescence intensity from the secondary-only control was subtracted from all samples (ΔMFI). Data are expressed as fold change in ΔMFI over the WT PGP1 control. Class-I HLA is stained with anti-HLA class-I antibody (EMR8-5 clone) (n = 3). *P*-values were calculated by two-way ANOVA *t*-test (**** *p*<0.0001). (**e**) Schematic showing the differentiation protocol of hPSC derived thymic epithelial like-cells. (**f**) Representative images of FlpOut ΔABC, FlpOut mini-1 and synHLA cell lines upon differentiation into hPSC derived thymic epithelial like-cells are shown in phase contrast (upper panel) (scale bar = 200 μm). Lower panel: FOXN1 (red) expression upon doxycycline induction (scale bar = 200 μm) **(g-h)** Expression of *FOXN1* and *DLL4* genes was measured by RT-qPCR in WT PGP1 (hPSC), FlpOut ΔABC, FlpOut mini- and synHLA cell lines carrying the inducible *FOXN1* circuit. Data represents mean ± SEM from n = 3 independent experiments. *P*-values were calculated by two-tailed Student’s *t*-test (\**p* < 0.05, **0.01, ***0.001). **(i)** Flow cytometry for anti-pan class-I HLA presentation into hPSC derived thymic epithelial like-cells (TEC-like cells) with and without IFN-γ generated from FlpOut ΔABC, FlpOut mini-1 and synHLA cell lines carrying the inducible *FOXN1* circuit. Data represent mean ± SEM from n = 3 independent experiments. *P*-values were calculated by unpaired two-tailed *t*-test (*p* < * 0.05, **0.01, *** 0.001, ns= not significant).

Collectively, these results demonstrate that the downregulation of the synthetic class-I HLA expression upon extended culture (P55) was not permanent genetic silencing or loss but rather reflects a cell fate dependent reconfiguration that mirrors native wild-type dynamics, confirming that both synthetic loci remained fully competent and responsive upon somatic differentiation.

Finally, we also examined whether engineered hPSCs retained the ability to differentiate into a developmentally relevant epithelial lineage in which HLAs are central ^49^. To confirm differentiation capacity of the engineered hPSCs, we forward programmed them toward a thymic epithelial cell (TEC)-like state using our previously validated doxycycline-inducible *FOXN1* genetic circuit site-specifically integrated at the ROGI1 locus by CRISPR-cas9 assisted HDR ^49^ into ΔABC, synHLA-, and mini-synHLA-engineered hPSCs (Fig. 6e). Differentiated cells exhibited increased expression of the inducible *FOXN1* (Fig.6f, 6g) and the canonical TEC marker *DLL4* (Fig. 6h), confirming lineage specification. Flow cytometry revealed basal class-I HLA expression in both mini- and full synHLA TEC-like cells and their high responsiveness to IFN-γ stimulation via elevated HLA expression (Fig. 6i).

These results confirm that large-scale genome restructuring using REWRITE does not impair differentiation potential, providing a path toward building HLA-specific somatic cells of interest from a common isogenic background.

## Discussion

Our work introduces REWRITE, a platform for scar-minimized rewriting of multigenic regions in human pluripotent stem cells (hPSCs). Although recent genome integration technologies have expanded programmable DNA insertion capabilities ^50^, genome engineering in hPSCs has remained largely limited to single-gene modifications or smaller payload integrations (<10 kb) ^31,32^. As a result, systematic refactoring of extended native human loci in situ — particularly in fragile hPSCs — has remained a major technical barrier.

By demonstrating >100 kb rewriting of the genetically complex HLA region ^6^, REWRITE establishes a practical benchmark for locus-scale rewriting of complex native genomic regions in hPSCs. Through integration and staging of established technologies, including CRISPR-mediated locus access and a unique RMCE “promoter-dock” architecture, REWRITE enables precise engineering of complex, repetitive loci in hPSCs while addressing practical constraints associated with payload delivery, size limits, and context-dependent silencing ^51–54^. While further advances will be required to extend payload size integration beyond current biophysical constraints ^41^, REWRITE provides a reproducible workflow for large-scale in situ genomic refactoring, enabling capabilities previously inaccessible in hPSCs ^19^.

Surprisingly, the integrated synthetic HLA architectures were initially highly active in hPSCs, which is counter to the expected low expression state of these sequences in hPSCs ^44^. While this state persisted for many passages, both HLA architectures eventually re-acquired the native low activity state, upon which the global transcriptional identity became near indistinguishable from parental wild-type hPSCs. Interestingly, while the full-length construct recapitulated the native locus transcriptional context by passage 55, the mini-synHLA resisted complete silencing at late passages with a remaining 45% expression level. Crucially, our RNA-seq data demonstrated that the enhanced transcriptional activity is confined to the synthetic loci only, without compromising the surrounding neighboring genes.

More importantly, the linear regression analysis of class-I HLA positive cell expression over time indicated that the passage-dependent decline for mini-synHLA follows the same, highly predictable trajectory as for full-length synHLA. They shared a similar coefficient of determination (R^2^=0.94) and the regression model suggested that mini-synHLA will reach the same stable baseline level of expression observed in WT hPSCs around passage 94.

Assessment of HLA proteins in the engineered cells via EMR8-5 and conformation specific W6/32 antibodies ^46–48^ demonstrated that the compact architecture of the mini-synHLA locus, similarly to the full-length synHLA architecture, allows for native transcription, translation, and proper trafficking of the proteins to the cell surface. Furthermore, the sustained activity of the locus does not cause an accumulation of misfolded proteins; on the contrary, the synthetic heavy chains are properly processed by the cells to form a properly-folded antigen-loaded HLA complex ^46–48^. Moreover, relative MFI analysis indicated that both engineered cell lines mirrored interferon responsiveness of the WT hPSCs.

Inclusive of our systematic characterization, this indicates that REWRITE enables generation of genomically stable hPSCs that converge toward a native transcriptional state while preserving key properties of pluripotent cells.

This is particularly important given the combinatorial diversity of class-I HLA haplotypes, where immune specificity arises from defined multi-allelic combinations rather than individual alleles in isolation^1^. Although the present study focused on establishing the engineering and regulatory tractability of rewritten HLA loci, future work will be needed to determine how specific synthetic haplotypes shape downstream immune interactions across differentiated cellular contexts and diverse donor backgrounds. By enabling installation of complete HLA haplotypes at native genomic sites, REWRITE provides a platform for systematic interrogation of HLA polymorphism and, more broadly, for programmable specification of complex genomic loci in an isogenic human pluripotent stem cell background.

## Acknowledgments

We would like to acknowledge members of the Neochromosome team for their contributions to assembling the synHLA variants.

## Author contributions

Conceptualization: DMT

Methodology: SFG, SL, SJ, MK, SD, SFP, BY, TY, TLD, NA, MH, DMT

Investigation: SFG, SL, SJ, MK, SD, SFP, BY, TY, TLD, NA, MH, DMT

Visualization: SFG, SL, SJ, MK, SD, SFP, BY, TY, DMT

Funding acquisition: DMT, LAM

Project administration: DMT

Supervision: DMT

Writing – original draft: SFG, SL, SJ, SD, SFP, TY, DMT

Writing – review & editing: SFG, DMT

## Data and materials availability

All raw fastq files from whole-genome sequencing and RNA-seq are available at BioProject PRJNA1327383. Plasmids and cell lines are available through a materials transfer agreements (MTAs) and will be made available via Addgene. All other data are available in the main text or the supplementary materials.

## Supplementary Materials

Materials and Methods

Supplementary Figs. S1 to S12

Supplementary Tables S1 to S5

