## Supplemental Files for "REWRITE Enables Locus-Scale Engineering of Synthetic HLA Haplotypes in Human Pluripotent Stem Cells"

**The PDF file includes:**

Materials and Methods
Supplementary Figs. S1 to S12
Tables S1 to S5

### Materials and Methods

hPSC culture and maintenance.

Human male induced pluripotent stem cells (iPSCs) PGP1 (obtained from George Church's laboratory) and female iPSCs NCRM2 lines (obtained from NINDS human cell repository) were maintained on Cultrex (R&D systems, Inc, MN, USA. RGF BME, Type 2, 3533-005-02) coated dishes at 37°C and 5% $CO_2$ in B8 medium [53] containing DMEM/F12 (Corning, NY, USA. 10092CM), 200 µg/mL of L-Ascorbic acid 2-phosphate (FUJIFILM Wako, 321-44823), 5 µg/mL of human insulin (Gibco, A11382IJ), 5 µg/mL of human transferrin (InVitria, 777TRF029), 20 ng/mL of sodium selenite (Thermo Fisher Scientific, CAS: 10102-18-8), 40 ng/mL of fibroblast growth factor 2-G3 (FGF2-G3) (made at Northwestern University's core facility), 0.1 ng/mL of neuregulin 1 (NRG1) (Peprotech, 100-03-1MG), and 0.1 ng/mL of transforming growth factor beta-3 (TGFb3) (Peprotech,10036E50UG) andpassaged using 0.5mM EDTA (NYU Reagent Preparation Facility). For maintenance, cells were passaged every 4–5 days. When single cell dissociation was required, cells were passaged with Accutase (Innovative Cell Technologies, NC9464543).

Cloning of landing pads and payload delivery vectors.

Landing pad (LP) and payload delivery (PV) vector plasmids were cloned using a combination of Gibson cloning, restriction digest cloning, and yeast recombineering using both naturally derived and chemically synthesized DNA sequences. The payload vector (FlpOut) are based on an arabinose copy-number inducible Bacterial Artificial Chromosome (BAC) and Yeast Artificial Chromosome (YAC) vector backbone. The landing pad plasmids contain four BsaI cut sites for inserting homology arms at the 5'- and 3'-ends of the cassette using Golden Gate cloning. Due to methylation restriction of two of these BsaI sites, propagation of the payload vector plasmids must be in *E. coli* strains lacking dam methylation. Payloads within payload vectors with the BAC markers must be propagated in a copy-number inducible *E. coli* strain such as EPI300 (Lucigen, EC300110).

Development of MAD7 plasmid.

The plasmid construct used to express erCas12a (pTL1) was based on the Cas9 plasmid pX459 available from Addgene (#62988). This is an all-in-one construct containing a U6 promoter-driven gRNA, a CbH/CAG promoter-driven Cas9 linked to a puromycin resistance cassette via a ribosome skipping P2A sequence. Mammalian codon-optimized Cas12a was commercially synthesized and cloned in place of the Cas9 gene. The Cas12a gRNA was cloned in place of the Cas9 gRNA. The direct repeat was modified at the polyT tract to contain two G nucleotides as suggested by Inscripta. The plasmid uses BsaI sites for golden gate cloning of new gRNAs. Guide sequences were determined using the CRISPOR database based on the algorithm for Cpf1 called DeepCpf1.

Landing Pad integrations.

Integration sites were identified based on suitable Cas9 gRNA locations. Cas9 gRNAs were identified for each target locus using the UCSC Genome Browser CRISPOR database, and the chosen guide sequences were assembled via Golden Gate Assembly into plasmid pX459. Sequences were chose based on high specificity (>70) and high activity scores (>65). To target the landing pad to these sites, short homology arms (500–800 bp) corresponding to the site were amplified from iPSC genomic DNA using primers designed to append Golden Gate compatible BsaI cut sites. Golden gate assembly was used to clone amplified homology arms into the landing pad plasmid and confirmed via colony PCR and sanger sequencing.

To target the genome, the landing pad was linearized in vitro using unique restriction enzymes NheI and KpnI and purified via Zymo's gel extraction kit protocol (D4007/D4008) prior to nucleofection. Human iPSCs were cultured to 60–80% confluency (~1.0 x $10^6$) cells and then treated for 2 hours with Thiazovivin ROCK inhibitor (Cayman Chemical, 14245). Cells were gently dissociated to a single cell suspension using accutase and then were nucleofected with 500 ng of the linearized landing pad, 1 µg of each CRISPR guide plasmid, and 1 µg *BCL-XL* plasmid using the Lonza P3 Primary Cell 4D-nucleofector kit (V4XP-3024) and CB150 program for hESCs, and plated into B8 with Thiazovivin (Cayman Chemical Company, 14245), and 20 µM Nedisertib/M3814 to inhibit NHEJ. Fresh B8 media was changed the day after nucleofection, and cells were allowed to recover for 2–4 days before starting selection with 3 µg/ml Blasticidin (Gibco A11139-03). After 5–7 days of selection, colonies were typically ready for manual clone picking. Colonies were imaged to confirm red fluorescence from the landing pad. Picked colonies were replated into a 96 well plate and expanded until filling a 6 well plate, then frozen down. PCR on genomic DNA was used to confirm correct integration of the landing pad in the desired location.

<u>Scarless biallelic deletion of HLA-A in human iPSCs.</u>
To achieve scarless biallelic deletion of the HLA-A locus, we utilized a combination of single-stranded oligonucleotide (ssODN) donor DNA and transiently expressed Mad7 nuclease to facilitate homology-directed repair. We designed two Mad7 gRNA pairs targeting the upstream and downstream sequences of an 8 kb region spanning HLA-A. Each gRNA was cloned by Golden Gate assembly into the puromycin-resistant plasmid pTL1. Four corresponding ssODNs were designed for the four gRNA combinations, each containing 50 nucleotides of homology on each side (100 nt total) and stabilized with two phosphorothioate bonds at both the 5'- and 3'-ends. For the deletions, 1 x 10^6 PGP1 hPSCs at 60–80% confluency were nucleofected with a pair of gRNA plasmids (2 µg each) and the corresponding ssODN (10 µg), supplemented with ROCK inhibitor Thiazovivin, Alt-R HDR enhancer V1 (IDT), and a BCL-XL expression plasmid. Following nucleofection, cells were replated in hPSC media with ROCK inhibitor for 24 hours. The media was then exchanged for hPSC media containing 0.3 µg/ml puromycin (Gibco, A11138-03) for a 48-hour selection period. After selection, cells were grown in standard hPSC media for an additional 10 days, or until colonies were visible. Individual colonies were manually picked, transferred to a 96-well plate for expansion, and screened by PCR for biallelic deletion. One gRNA/ssODN combination yielded the final deletion clone, which was validated by PCR and whole-genome sequencing (WGS).

<u>Assembly of synHLA and mini-synHLA.</u>
The synHLA and mini-synHLA constructs were assembled using yeast recombineering, with genomic DNA fragments sourced from BACs RP11-192H11 and CH17-93C5 (BACPAC Resource Center).

<u>mini-synHLA (38 kb).</u> The compact construct was assembled in a single step in S. cerevisiae (strain BY4741). Six PCR-amplified fragments (4–5 kb each) corresponding to the genomic loci of HLA-A, -B, and -C were co-transformed with 50 ng of linearized FlpOut vector. The fragments included predicted enhancers and native polyadenylation signals. Synthetic DNA linkers with terminal homology were used to bridge the fragments. Following transformation, yeast were

selected on SC-His dropout media. Correct assemblies were identified by colony PCR, and the plasmid DNA was extracted from yeast, electroporated into EPI300 E. coli for amplification, and sequence-verified by Illumina deep sequencing (fig. S5).

synHLA (115 kb). The full-length construct was assembled in a two-step yeast recombineering process. First, the 93 kb region containing the *HLA-B* and *HLA-C* loci (hg38 chr6:31267251–31359518) was captured from BAC CH17-93C5 into the FlpOut vector. Second, this intermediate plasmid was linearized in vitro with a site-specific CRISPR/Cas9 RNP (targeting chr6:31315895). The linearized plasmid was then transformed back into yeast along with two 4 kb PCR fragments covering the *HLA-A* locus, which were inserted via a second round of recombineering. The final, fully assembled synHLA plasmid was recovered, amplified in E. coli, and validated by Illumina deep sequencing as described above (fig. S5).

Assembly of synHLA allelic variants.

To generate the allelic variants in the full synHLA construct, we utilized a two-step process. To avoid shear stress on synHLA in vitro given its size and repetitiveness, we used a yeast recombineering approach utilizing positive/negative selection of *URA3*, which may be positively selected using uracil drop-out media and negatively selected using 5-fluoroorotic acid (5-FOA; Gold Biotechnology). To swap specific alleles, the *URA3* gene was engineered in place of polymorphic exons 2 and 3 of the intended HLA through yeast homology-directed recombination (HR) (fig. S5C). Allelic HLA-variants of these exonic regions were chemically synthesized and used as DNA-repair donors for CRISPR/Cas9-assisted HR in place of *URA3*, followed by counter-selection with 5-FOA revealing yeast colonies with precise allelic replacement (fig. S5C). Colonies were screened using high-throughput PCR. Correct constructs were transferred to EPI300 *E. coli* (Lucigen, EC300110) after spheroplasting and alkaline lysis precipitation followed by electroporation. Constructs were verified using deep sequencing.

Assembly of mini-synHLA allelic variants (haplotypes).

Individual HLA allele/regulatory sequence plasmids were cloned via yeast recombineering into the yeast pWS171 backbone using custom DNA fragments (IDT) to connect the regulatory fragments to the backbone. Colony PCR was performed on yeast to confirm correct plasmid assembly. Yeast plasmids were extracted and transformed into *E. coli*, and extracted plasmid was further analyzed by restriction digestion and then whole plasmid sequencing (Azenta). Unique restriction sites were added to exons 1 and 8 for swapping out HLA alleles. New HLA alleles were synthesized by either Qinglan (China) or Twist Biosciences (USA) and then cloned in place of the exons of the parental construct. Desired HLA-A, -B, and -C allele combinations were digested and assembled with digested FlpOut plasmid backbone to create a new mini-synHLA haplotype plasmids. Correct mini-synHLA assembly was confirmed via yeast colony PCR on four assembly junctions, and subsequently with restriction digestion and whole plasmid sequencing.

Assembly of 62kb construct.

For intermediate-size benchmarking, a 62 kb payload was assembled from a contiguous human genomic sequence adjacent to the HLA class-I region (hg38 coordinates chr6:31,260,000–31,360,000). This construct was used exclusively for evaluating size-dependent integration efficiency and was not functionally characterized. The 62 kb construct was assembled in a single step in *S. cerevisiae* (strain BY4741). Human genomic DNA was isolated from NCRM2 cells and used as a template to amplify 14 PCR fragments (4–7 kb each corresponding to the genomic locus, with terminal overlaps ~100 bp in length on average. Primer and linkers sequences are listed in Supplmentary Tables S1 and S2. They were co-transformed with 100 ng of linearized payload vector, Zero1, containing the PuroR and mNeonGreen genes. The left and the right linked fragments to the vector were synthetized by Twist Biosciences (USA). Primer sequences are listed in Supplementary Table S1. Following transformation, yeasts were selected on SC-His dropout media. Correct assemblies were identified by colony PCR, and the plasmid DNA was extracted from yeast, electroporated into EPI300 *E. coli* for amplification, and sequence-verified by nanopore sequencing.

Payload DNA deliveries.

Human iPSCs containing the REWRITE landing pad (LPcore) were passaged via EDTA two days before nucleofection, to achieve a confluency of 60–80% on the day of nucleofection. Rock inhibitor was added to the cells 2–4 hours prior to nucleofection and then dissociated to a single-cell suspension using accutase. The number of nucleofected cells scaled with the size of the payload DNA, with amounts ranging from as a low as 2 x $10^5$, 1 x $10^6$, and 5 x $10^6$ cells. Nucleofection was done using Lonza P3 Primary Cell 4D-nucleofector kit using the CB150 program for hESCs and allowed to recover overnight in B8 with rock inhibitor thiazovivin. Each delivery used 5–10 µg payload plasmid, 2 µg pCAG-iCre (Addgene #89573), and 2 µg BCL-XL plasmid. The morning after nucleofection, the media was changed to B8 with no rock inhibitor. Cells were allowed to recover for an additional 3–5 days. Selection for integration of the payload used 0.3 µg/ml puromycin for FlpOut deliveries. Colonies surviving selection were picked after 7–10 days of selection into a 96 well plate. After 2–3 days, picked colonies were passaged via EDTA into a new well of the 96 well plate, and 10 µl of cell suspension was saved for digestion with proteinase K for colony PCR. Colonies were imaged and observed for the presence of mNeonGreen or mScarlet. Colony PCR was performed on extracted genomic DNA to verify the integration of the payload plasmid into the landing pad. Colonies with confirmed payload delivery and expected morphology were clonally expanded.

FlpOut induction marker excision.

Cells containing the FlpOut cassette were passaged and maintained in puromycin to ensure expression of thymidine kinase. Before starting the excision, cells were passaged via accutase and seeded into a 12 well plate at low densities per well. Flp recombinase was induced by adding 200 nM 4-hydroxytamoxifin (4-OHT; Sigma Aldrich, 68392-35-8) during the replating. Cells were exposed to 4-OHT for 24–72 hours, after which successful cassette excision was selected for using 500 nM ganviclovir against the presence of thymidine kinase in the media. After 6 days under ganciclovir selection, surviving colonies were picked and screened for successful excision of the FlpOut cassette. PCR was used to confirm the removal of the cassette, and a portion of post-excision cells were placed under puromycin selection to confirm lack of resistance.

RT-qPCR assays.

RNA was extracted using the Quick-RNA Miniprep Kit (Zymo Research, CA, US, R1054/R1055) and quantified by NanoDrop spectrophotometer (Thermo Fisher Scientific, MA, USA). cDNA synthesis was performed using ABScript Neo RT Master Mix (ABclonal, Inc., MA, USA, RK20432). RT-qPCR reactions were set up in triplicate with the Forget-Me-Not EvaGreen qPCR Master Mix manufactured by Biotium (CA, USA) (Fisher, NC1481635). Reactions were run on a

QuantStudio-5 Real-Time PCR System (Thermo Fisher Scientific, MA, USA) with 40 cycles of 30 s at 95°C, 30 s at 60°C and 30 s at 72°C. An endogenous control, the human RSP29 gene, was used to normalize the expression. The relative gene-expression levels were calculated by the 2-ΔΔCt method. Primers are provided in Table S4.

Alkaline phosphatase assay.
Alkaline phosphatase assays were performed using the Alkaline Phosphatase Staining Kit II (Stemgent) following the instructions provided by the manufacturer (Stemgent, Amsbio #AMS.00-0055).

Endothelial cell differentiation.
Endothelial cell differentiation from human iPSCs was performed based on the adaptation of the protocol from [54]. Human iPSCs at 80% confluency were differentiated into mesoderm cells using RPMI (Corning, 10-011-CV) supplemented with B-27 supplement minus insulin (Gibco, A18956-01) and 6 μM CHIR99021 (Stemcell technologies, 72054) for two days. After mesoderm induction, media was changed to EGM-2 medium (Lonza, CC-3162) supplemented with 50 ng/mL VEGF (Sino Biological, 10008-HNAH) and 25 ng/mL FGF2 (Sino Biological, 10014-HNAE) for 3 days. Cells were split and passaged with TrypLE (Gibco, 12605010) onto gelatin coated plates and maintained in EGM-2 supplemented with VEGF and FGF-2 and were maintained by passaging 1:2 with TrypLE Express when confluency became high.

Induction of thymic epithelial cells.
To generate thymic epithelial cells, we first introduced a doxycycline-inducible *FOXN1* circuit, previously developed and reported by our group, into the ROGI1 locus of HLA-edited iPSCs using nucleofection [46]. The thymic epithelial cell differentiation was performed by adapting the protocol from [55]. Once the modified iPSCs reached approximately 40–50% confluency, the culture medium was replaced with differentiation medium composed by Advanced DMEM supplemented with 60 mg/L 2-ascorbic acid 2-phosphate (FUJIFILM Wako, 321-44823), 10 mL/L GlutaMAX (Gibco, 35050-61) and 15 mL/L 1 M HEPES (Gibco, 25-061-CI) and the addition of different cytokines at different time points: (day 1 to day 4: Activin A (Sino Biological, 10429-HNAH), WNT3A (MyBioSouse, MBS1265206), FGF2 (Sino Biological, 103623-738), and TTNPB (AmBeed, A700949); from day 5 to day 8: BMP4 (Peprotech, AF12005ET10) TTNPB (AmBeed, A700949), LY364947 (MedChem express, HTS466284); from day 9 to day 12: BMP4 (Peprotech, AF12005ET10), WNT3A (MyBioSouse, MBS1265206), FGF8b (Sino Biological, 16277-HNAE), TTNPB (AmBeed, A700949) and LY364947 (MedChem express, HTS466284)) FOXN1 genetic circuit was induced by the addition of doxycycline (dox) (Sigma-Aldrich, D5207) at a final concentration of 100 ng/mL between day 5 and day 8 At day 12 the cells were collected to perform either RT-qPCR or proceed with IFN-γ stimulation for the downstream flow cytometry analysis.

Immunostaining.
For imaging, cells were grown directly on coverslips (Fisher Scientific, 12-541-055). They were fixed with PFA 4% (Alfa Aesar, 43368) for 10 min at RT and rinsed three times with PBS for 5 min each. Fixed cells were permeabilized in 0.5% Triton X-100 (EMS, 22140) in PBS buffer for 5 min at room temperature. Cells were blocked in blocking buffer (3% BSA, 0.01% Triton X-100 in PBS) (BSA, GoldBio, a421-100) for one hour at room temperature. Cells were incubated with primary antibody in blocking buffer overnight at 4C. Cells were washed three times for 5 min each

with wash buffer (2% BSA, 0.01% Triton X-100 in PBS) and incubated with the secondary antibody in blocking buffer for 1h at RT. Cells were washed three times for 5 min each with wash buffer. After rinsing, coverslips were mounted VECTASHIELD mounting medium with DAPI (H-1200-10) and imaged using an EVOS M7000 imaging system (60x oil objective). Primary and secondary antibodies are provided in the Supplementary Table S5.

<u>Flow cytometry.</u>
Cells were dissociated using Accutase for 5 min at 37°C and quenched with DMEM/F12. The cell suspension was centrifuged at 200 *g* for 5 min and resuspended in PBS with 2% Fetal Bovine Calf Serum (FCS) (Corning, 35-054-CM). Cells were stained with primary antibodies for 30 min at 4°C in the dark, diluted according to recommended dilutions. Following primary staining, cells were washed twice in PBS with 2% FBS. Secondary antibodies, when used, were diluted 1:500, and cells were stained for 30 min at 4°C in the dark. Cells were washed twice and stained with DAPI (1:2000, BD Pharmingen, 564907) for viability prior to a final wash and resuspension in PBS with 2% FBS. Unstained cells and secondary-only were used as negative staining control. Flow cytometry analysis was performed with a SONY SA3800 analyzer. Gates were adjusted on non-stained controls. In total, 10,000 events (alive cells) per sample were acquired. The data were plotted using FlowJo v10.8 Software (BD Life Sciences). Median Fluorescence Intensity (ΔMFI) was calculated by subtracting the MFI of the secondary-only control from the MFI of each stained sample. Relative expression levels were determined by normalizing the ΔMFI of each sample to the ΔMFI of the WT PGP1 control.

<u>IFN-γ treatment.</u>
The engineered human iPSCs lines (PGP1-ΔABC, mini-synHLA and synHLA) and the control PGP1 cell line were cultured for 2 days with or without recombinant human IFN-γ (50–100 ng/mL, Sino Biological, 11725-HNAS-100) for the assessment of HLA class-I expression. Upon treatment with IFN-γ, the cells were harvested and stained using specific monoclonal antibodies against pan HLA class-I (ABC) and analyzed with a SONY SA3800 flow cytometer in accordance with the manufacturer's instructions. Gates were adjusted on non-stained controls. In total, 10,000 events (alive cells) per sample were acquired. The data were plotted using FlowJo v10.8 Software (BD Life Sciences).

<u>Whole-genome sequencing.</u>
Genomic DNA was extracted following manufacturer’s instructions (Zymo Research, D3025) and submitted to Novogene for paired-end whole genome sequencing (150 bp paired-end reads, 30x depth, 90 Gb) on an Illumina NovaSeq X system. The returned raw FASTQ files were assessed using FastQC and MultiQC, then trimmed using Trimmomatic. Trimmed reads were aligned to the GRCh38/hg38 reference genome using BWA-MEM, followed by BAM conversion with Samtools, and deduplicated with Picard. Reference genome files were prepared by indexing the hg38 FASTA with BWA and Samtools and generating a sequence dictionary using Picard. Raw data is available at BioProject PRJNA1327383. To verify successful genome rewriting, BAM files were visualized on IGV to confirm removal of the HLA loci and replacement with the landing pad or synHLA. Circos plots were generated in R-studio using the package "Circlize". Variant calling was performed on BAMs with GATK and resulting VCFs were annotated using SnpEff and filtered with SnpSift to reference the Clinvar and SNPedia pathogenicity data. Variants were manually interpreted to ensure no pathogenic *TP53* or other high-impact mutations were present

(fig. S8G).

To verify successful deletion and/or integration, FASTQ files were aligned to a custom reference of hg38 containing the expected combinations of sequences of the LP, HygroTK, mini-synHLA, and full-synHLA cassettes as separate sequences from the native loci. BAM files were sorted with SAMtools and converted to bigWigs with bin sizes of 2000 bp. bigWigs were visualized on IGV to confirm removal of the HLA loci at expected locations in comparison to controls. Integrations sorted BAM files were checked for chimeric reads spanning the ends of the integration and the native chr6 locus and visualized.

The large language models (LLMs) ChatGPT, Claude Opus 4.1, and Perplexity AI Pro were used for assistance with bioinformatics pipeline development and debugging.

PGP1 parental genome HLA haplotype was assessed from BioProject PRJNA259786, by accessing SRA SRR1583555 and assessing the alignment on chromosome 6 to the GRCh38/hg38 reference genome.

RNA-sequencing.

Total RNA was extracted following manufacturer's instructions (Zymo Research, R1055) and submitted to Plasmidsaur for poly(A) enriched sequencing (150 bp single-end reads, 6 Gb per sample) on an Illumina NovaSeq X system. Raw data is available at BioProject PRJNA1327383. The returned raw FASTQ files were assessed using FastQC, trimmed for adapter sequences using Trim Galore, and "pseudo"-aligned with Kallisto to a Kallisto index generated from a custom reference built by removing all *HLA-A*, *HLA-B*, and *HLA-C* cDNA sequences within the hg38 reference. GRCh38.cdna.all.fa from ensemble and replacing them with haplotype specific cDNA sequences expected from our integrations or the native haplotypes of the PGP1 control.

Expression values were determined in R by converting Kallisto abundance data from transcript-level quantification to gene-level quantification using tximport. Gene level count data was converted to a dds object with DESeq2, and the subsequent differential expression results had lfcshrink with type = 'apeglm' performed on them. Heatmaps were generated from the differential expression analysis results using the pheatmap package in R. Due to the large variance between positive and negative pluripotency factors, expression values of pluripotency genes were visualized after a log2 transformation. To check haplotype specific alignment, Pseudobams were sorted using SAMtools and aligned to their custom references on IGV.

The large language models (LLMs) ChatGPT, Claude Opus 4.1, and Perplexity AI Pro were used for assistance with bioinformatics pipeline development and debugging.

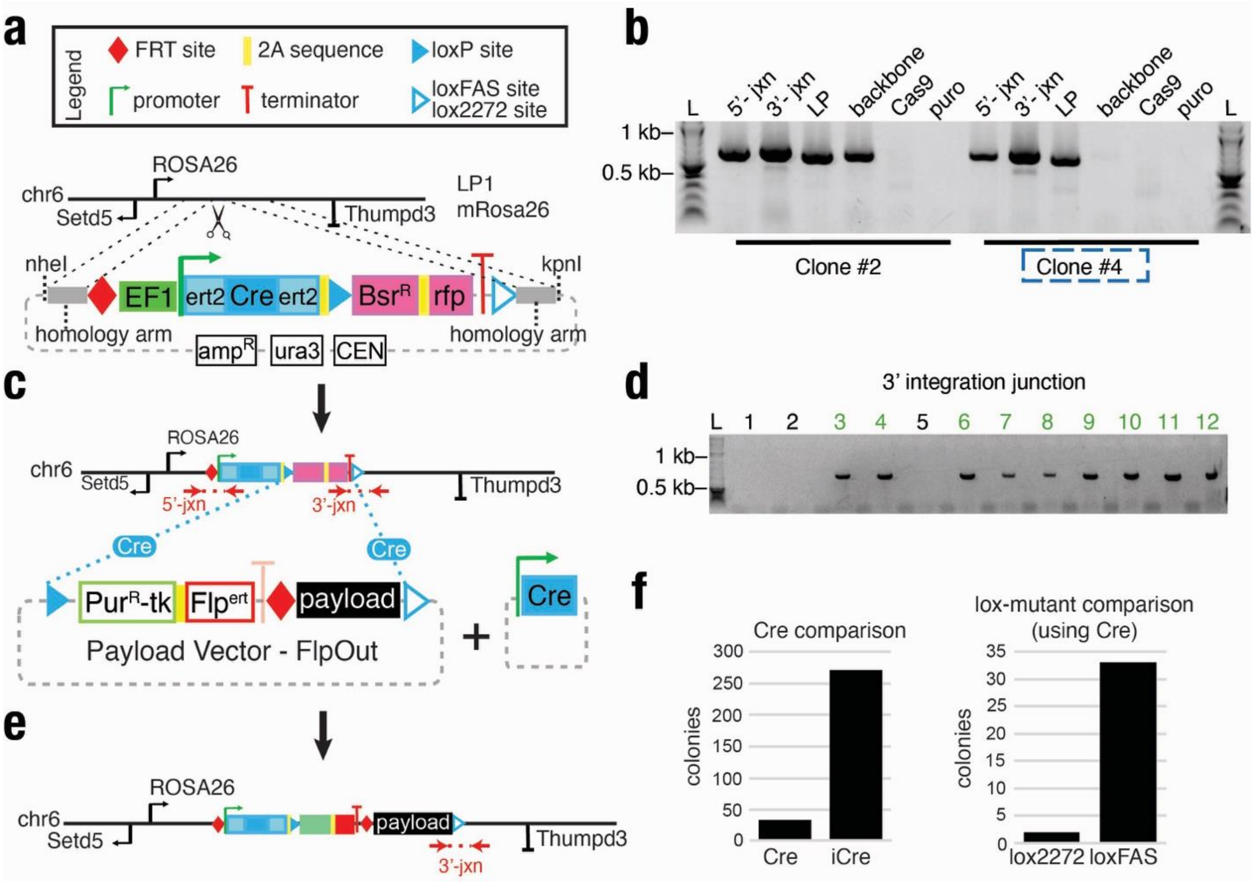


**Supplementary Figure S1. Benchmarking landing pad design, Cre activity, and RMCE efficiency in mESCs.**

**(a)** Schematic of LP1 integration into the ROSA26 locus in mouse embryonic stem cells (mESCs). The cassette includes a tamoxifen-inducible ERT2-Cre-ERT2 (double-flanked to reduce leaky expression), blasticidin resistance ($Bsr^R$), and mScarlet (RFP), flanked by wild-type loxP and loxFAS sites. Homology arms for CRISPR-assisted targeting flank the insert. Top key defines schematic elements.

(**b**) Junction (jxn) PCR validation of LP1 insertion in clones #2 and #4. Non-specific integration controls include backbone, Cas9-only, and puromycin vector.

(**c**) Schematic of Cre-mediated recombinase cassette exchange (RMCE) to replace LP1 with a terminal payload vector encoding a Puromycin–thymidine kinase ($Pur^R$–TK) fusion, FlpERT2, and a 10 kb fragment of human *HLA-A*.

(**d**) PCR screening showing 9 of 12 subclones had 3′ integration of payload.

(**e**) Final integrated configuration after RMCE showing scar-minimized replacement of the landing pad with payload vector.

(**f**) Comparison of recombination frequencies (left histogram), iCre yields significantly more colonies than Cre; right histogram: loxFAS outperforms lox2272 for RMCE using Cre.

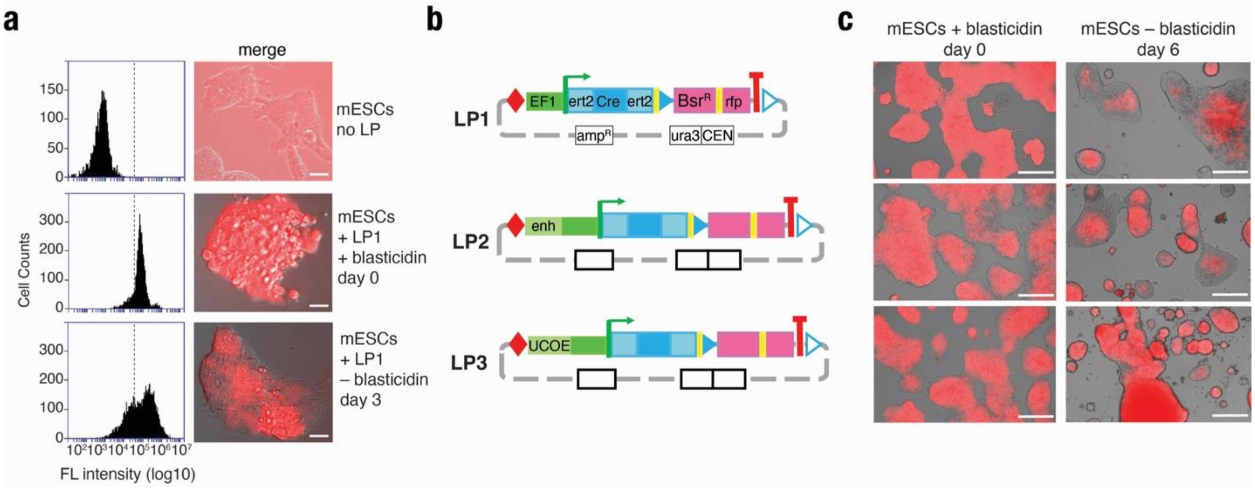


**Supplementary Figure S2. Universal chromatin opening element reduces REWRITE landing pad silencing.**

Evaluation of REWRITE landing pad expression in mESCs with or without chromatin-opening elements. LP1 (top row) includes an EF1α-driven ERT2-Cre-ERT2 cassette, blasticidin resistance (BsrR), and mScarlet (RFP). LP2 adds a synthetic enhancer upstream of EF1α. LP3 includes a Universal Chromatin Opening Element (UCOE) in place of the enhancer.

**(a)** Flow cytometry assay shows RFP intensity in mESCs with and without blasticidin selection. (Scale bars = 200 μm).

(**b**) Schematics of LP1–LP3 architectures highlighting promoter, enhancer or UCOE, recombinase, and selection elements.

(**c**) Fluorescence microscopy shows persistent LP silencing in LP1, partial silencing in LP2, and stable expression in LP3 after 6 days without blasticidin (Scale bar = 200 μm).

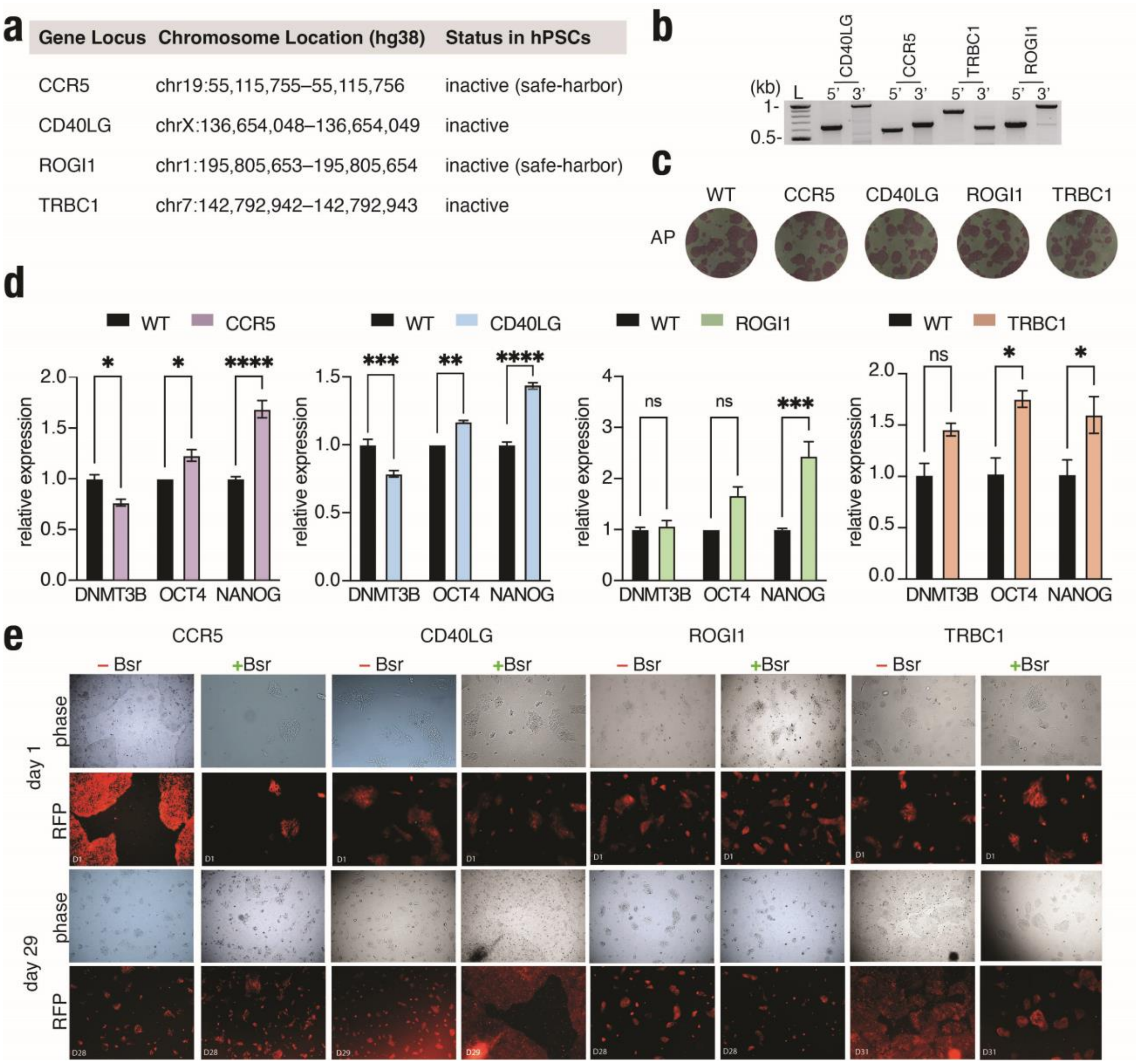


**Supplementary Fig. S3. Evaluation of REWRITE landing pad expression stability across genomic loci in hPSCs.**

**(a)** Table summarizing genomic loci targeted for LPneo integration in WT PGP1 hPSCs, including chromosomal location and transcriptional status in the undifferentiated state.

**(b)** Junction PCR validation of LPneo insertion at the different genomic loci.

**(c)** Images representing alkaline phosphatase staining of hPSCs upon LPneo integration at each locus. WT PGP1 as a control.

**(d)** RT-qPCR for pluripotency-associated genes (*DNMT3B*, *OCT4*, and *NANOG*) across hPSC clones with LPneo integrated at each locus. Gene expression is normalized to WT PGP1. Data shown as mean ± SEM (n = 3); *P*-values were calculated by two-way ANOVA ***t***-test (* ***p***<0.05, **<0.01, ***<0.001, ****<0.0001, ns = not significant).

(**e**) Fluorescence microscopy of RFP expression in hPSCs at day 1 and day 29 after LPneo integration at each locus, with and without blasticidin (Bsr) selection. Minimal silencing of RFP signal was observed at all loci over 29 days, even without selection.

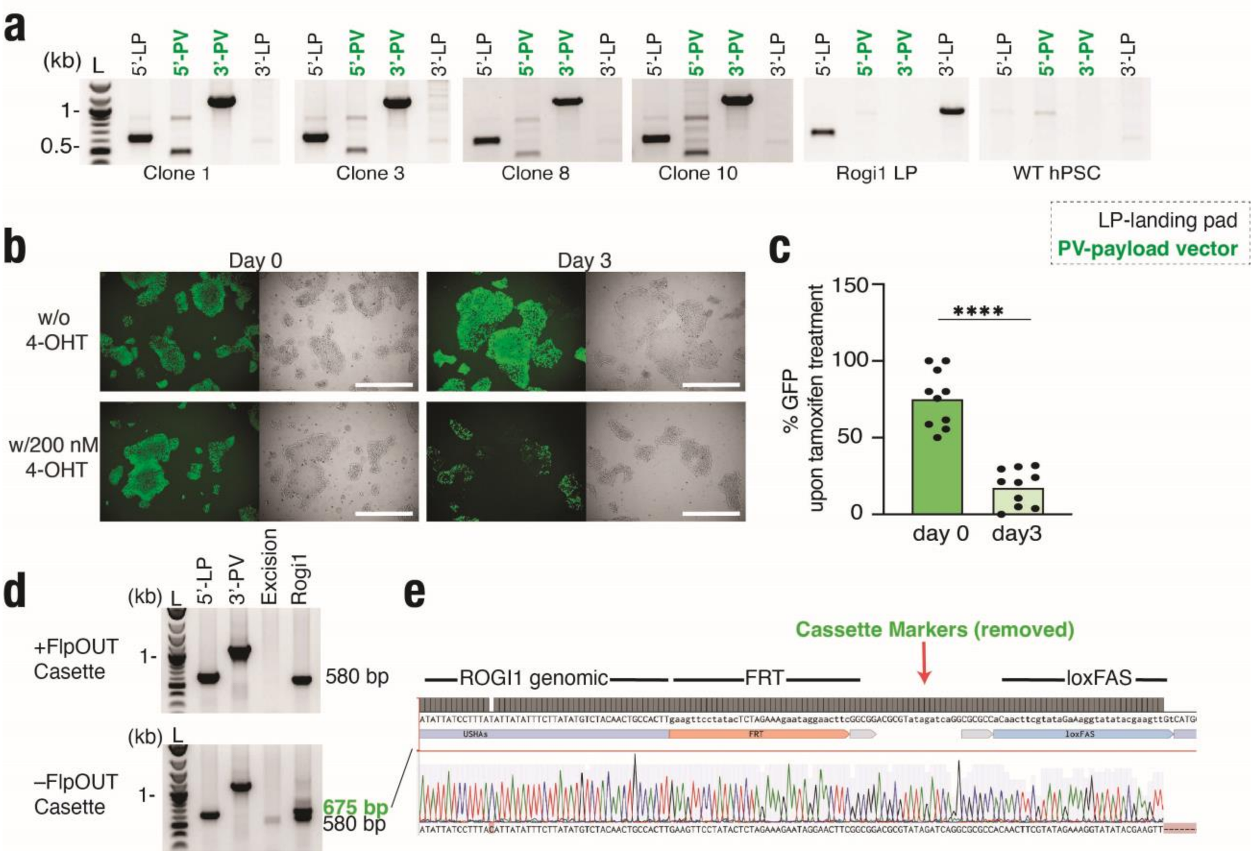


**Supplementary Figure S4. Tamoxifen induced excision of the FlpOut cassette at the ROGI1 locus**

**(a)** Junction PCR confirmation of FlpOut payload vector (PV) integration into the landing pad (LP).

**(b)** Tamoxifen induction of FlpERT2 kinetics at ROGI1 locus, as shown by GFP expression levels before and after induction for 3 days (*left*), and as phase contrast images (*right*). (scale bar = 200 μm)

**(c)** Average percent of GFP-positive cells within 10 FlpOut hPSC colonies at ROGI1 locus, calculated by counting cells in each colony (average cells per colony: day 0 =169.2, day 3= 381.6) before and after 3 days of tamoxifen exposure. *P*-value was calculated by paired *t*-test (****$p$<0.0001)

**(d)** PCR gel showing before and after excision of the FlpOut cassette. Excision allows for the amplification of a band between loxFAS and the upstream ROGI1 homology arm of 657 bp.

**(e)** Sanger sequencing confirms the excision of the FlpOut cassette.

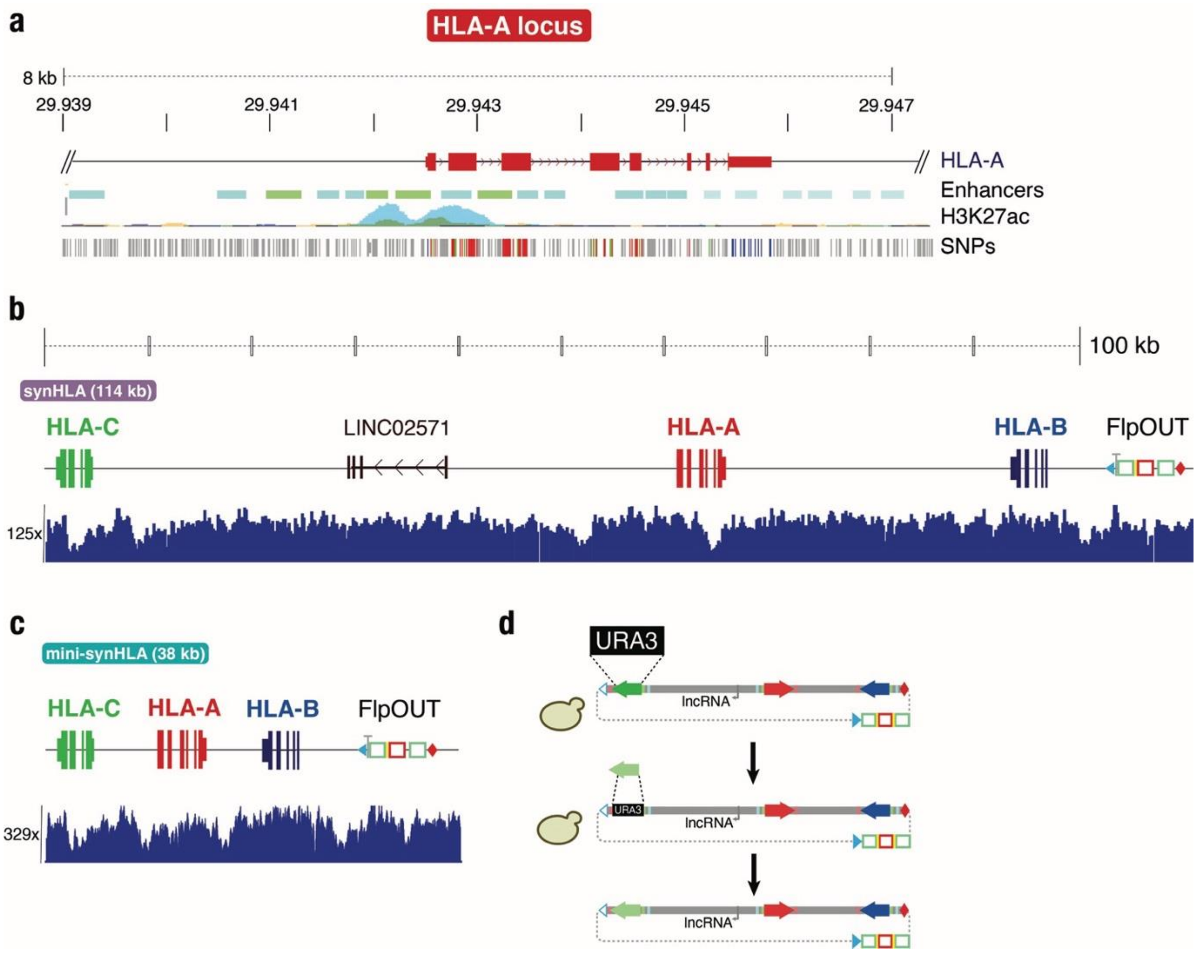


**Supplementary Figure S5. Validation of synHLA and mini-synHLA payload assemblies and recombineering strategy.**

**(a)** *HLA-A* locus, which was relocated within the *HLA-B* and *HLA-C* intergenic region. Tracks below show locations of ENCODE-predicted enhancers, which were used to define the "core locus" of ~8 kb. H3K27 acetylation track shows active marks in cells that express *HLA-A*, HUVEC (endothelial cells; light blue) and HSMM (skeletal muscle; green). However, there were no marks for H1-hESC (human embryonic stem cells; gold) consistent with lack of expression in pluripotent stem cells. SNPs (single nucleotide polymorphisms) track shows density of allelic variations in exons 2 and 3. As the class-I HLAs are homologs of each other, the regulatory structure shown here is similar for all class-I HLAs.
**(b)** Genome-wide sequencing coverage of the full-length 115 kb synHLA construct following recombineering and integration into human pluripotent stem cells. Coverage is uniform across the *HLA-C*, LINC02571, *HLA-A*, and *HLA-B* regions, confirming structural integrity and full-length assembly.
**(c)** Sequencing coverage of the compact 38 kb mini-synHLA construct, showing similarly uniform depth across all three class-I HLA genes.
**(d)** Schematic of yeast-based allele-swapping and recombineering workflow. The synHLA backbone is assembled in *S. cerevisiae* with allele-specific HLA fragments and selectable *URA3* marker. CRISPR/Cas9-assisted counterselection enables modular allele replacement for rapid haplotype customization.

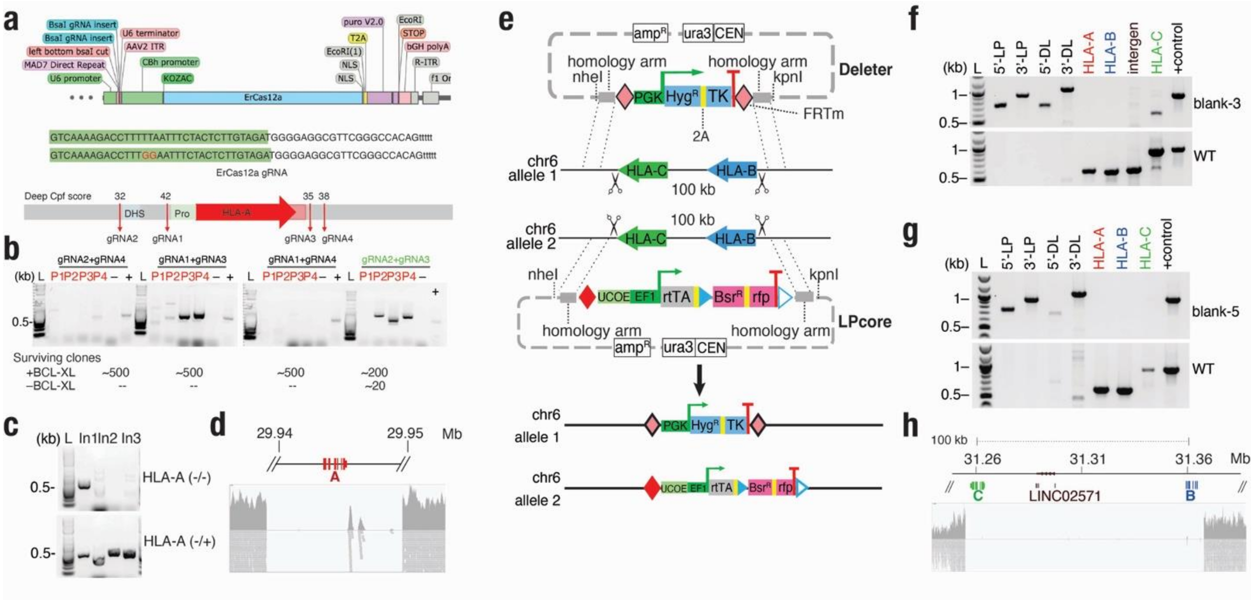

**Supplementary Figure S6. Biallelic deletion of *HLA-A* and generation of additional HLA-ABC deletion lines.**

**(a)** Design of Mad7 gRNAs for biallelic deletion of *HLA-A*.
**(b)** PCR screen of different Mad7 gRNA combinations for *HLA-A* deletion.
**(c)** PCR verification of biallelic deletion of *HLA-A* from WT PGP1 hPSC.
**(d)** Deletion of class-I *HLA-A* in WT PGP1 is shown via WGS against hg38.
**(e)** Schematic representation of CRISPR/Cas9 biallelic deletion of class-I *HLA-B/C* locus followed by LPcore and HygTK "deleter" cassette integration on chromosome 6.
**(f–g)** Junction PCR validation of LPcore and HygTK "deleter" (DL) cassette integration and class-I HLA genes deletion in different WT PGP1 hPSC clones (blank clone 3 and 5)
**(h)** Deletion of class-I *HLA-B/C* locus in WT PGP1 hPSC in blank-5 clone is shown via WGS against hg38.

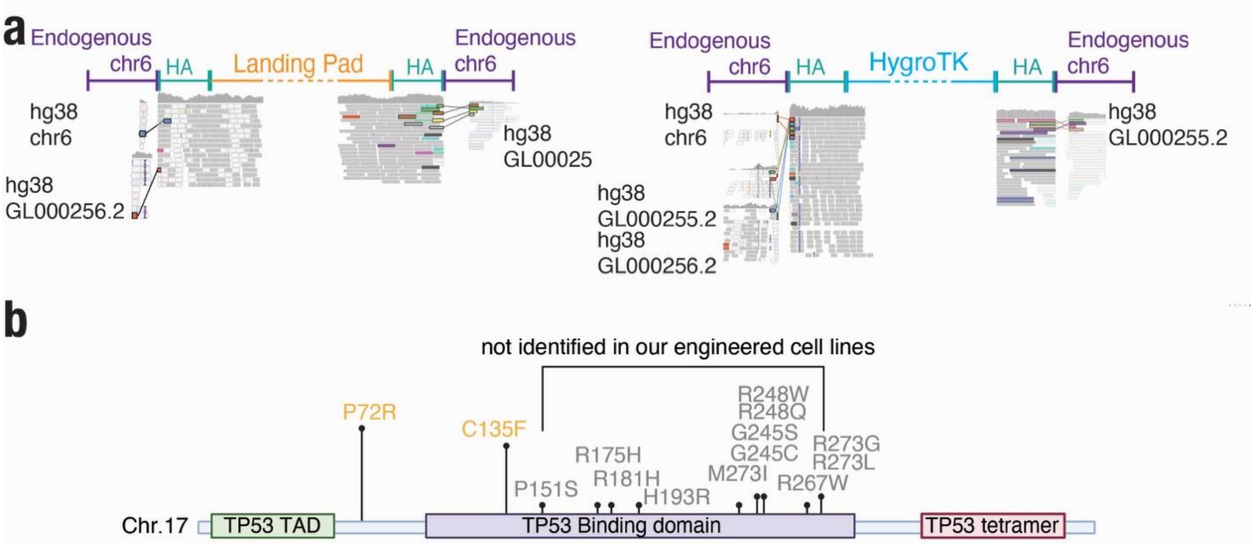


**Supplementary Figure S7. Verification of landing pad junctions at second region of chromosome 6.**

**(a)** Chimeric reads spanning landing pad (LPcore) (*left*) and HygroTK "deleter" (*right*) cassettes' homology arms for integration via Homology Directed Repair (HDR) and endogenous chr6 sequences and alternative sequences on both the 5'- and 3'-ends verifying integration via WGS. **(b)** Graphical representation of *TP53* gene. The most common mutations described in hPSCs and acquired with passage number under *in vitro* culture are in gray and they have not been identified in our engineered cell hPSC lines. In yellow, two polymorphisms are shown already present in the parental cell line (WT PGP1).

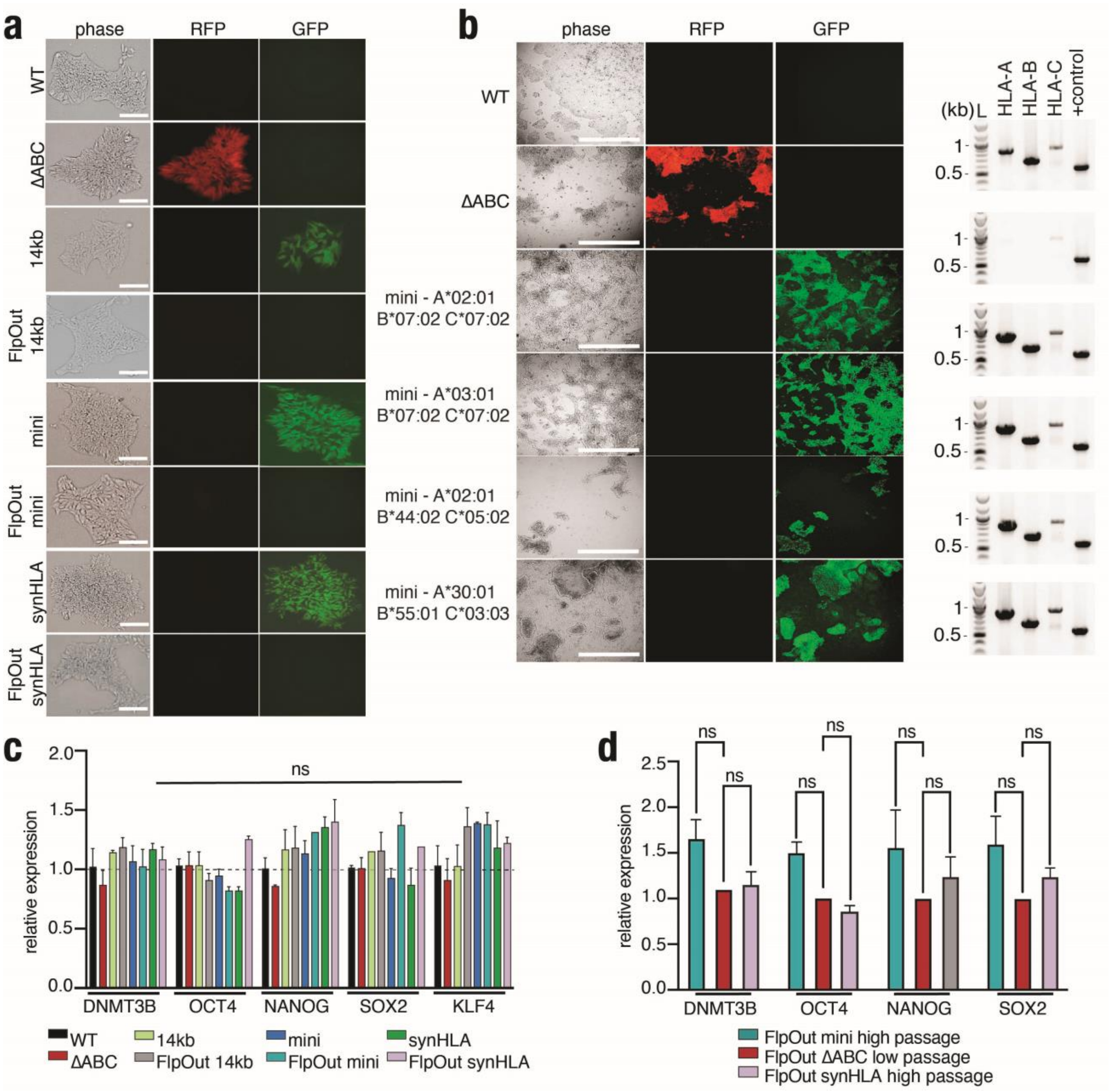


**Supplementary Figure S8. Synthetic HLA delivery in hPSCs and pluripotency verification.**
**(a)** Representative images of WT PGP1, ΔABC, mini- and synHLA integration before and after flipping out the cassette are shown in phase contrast, with the landing pad integration (RFP), and the payload deliveries (GFP). (Scale bar = 200 μm).
**(b)** Representative images of three different class-I HLA haplotypes assembled in mini synHLAs and delivered to the ΔABC clone (scale bar = 650 μm). PCR gel verifying the deletion in the ΔABC clone and the integration of class-I HLA genes upon delivery. WT PGP1 as a control.
**(c)** Expression of pluripotency genes was measured by RT-qPCR in WT PGP1, ΔABC, mini-1 and synHLA integration before and after flipping out the cassette. Gene expression was normalized to WT PGP1 (dotted line). Data represent mean ± SEM from n = 3 independent

experiments. *P*-values were calculated by two-way ANOVA t-test (* $p<0.05$, ns = not significant).

**(d)** RT-qPCR of pluripotency markers on high passage (p22). FlpOut mini-1 and synHLA compared to low passage (p7) ΔABC cells. Data represent mean ± SEM from n = 3 independent experiments. *P*-values were calculated by unpaired two-tailed *t*-test (* $p<0.05$, ns = not significant).

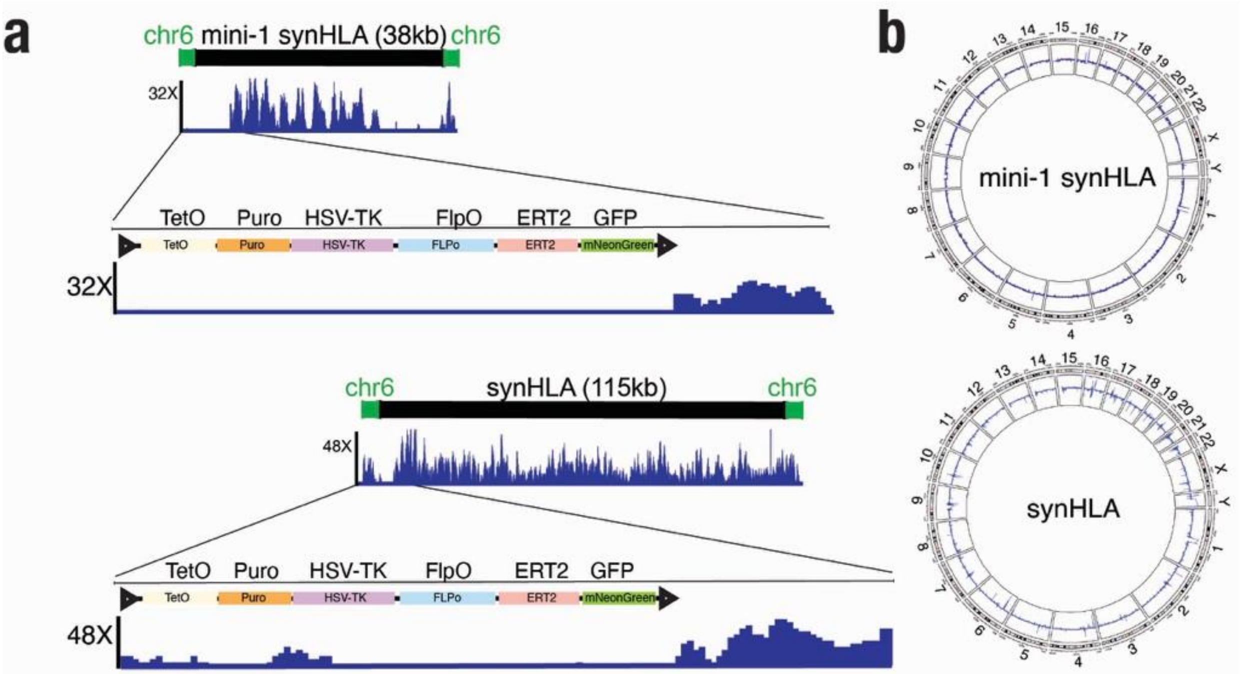


**Supplementary Figure S9. Verification of complete marker excision.**
**(a)** Whole genome sequencing of mini-1 and synHLA showing the absence of reads corresponding to marker cassettes comprising e.g., PuromycinR, mNeongreen, and Flp-ERT2, confirming complete excision.
**(b)** Circos plot showing no chromosomal aberrations of FlpOut mini-1 and synHLA hPSC. WT PGP1 as a control.

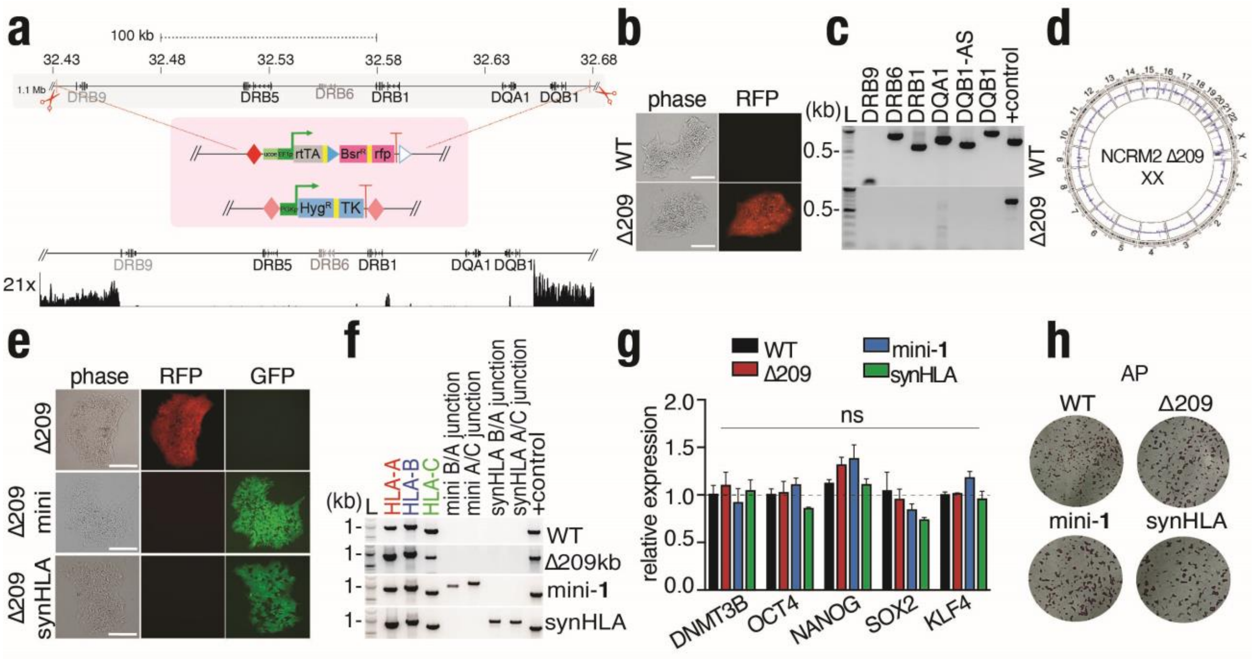


**Supplementary Figure S10. Integration of mini-1 and synHLA into the class-II HLA region in NCRM2 hPSC line.**

**(a)** CRISPR-based deletion of 209 kb class-II HLA cluster in female NCRM2 hPSCs.

**(b)** Representative images of WT NCRM2 and Δ209 kb class-II HLA hPSCs shown in phase contrast and with the landing pad integration (RFP). (Scale bar = 200 μm)

**(c)** PCR gel verifying *HLA-DRB1*, *-DQA1*, *-DQB1* class-II gene biallelic deletions in Δ209 kb hPSC. WT NCRM2 as a control.

**(d)** Circos plot showing no chromosomal aberrations of NCRM2-Δ209 hPSC.

**(e)** Representative images of mini-1 and synHLA integration in NCRM2-Δ209 shown in phase contrast, with the landing pad integration (RFP), and the payload deliveries (GFP). (Scale bar = 200 μm).

**(f)** PCR gel verifying the integration of the class-I HLA genes upon delivery of the mini-**1** and synHLA payloads. WT NCRM2 as a control. Specific junctions between *HLA-B/A* and *HLA-A/C* are amplified to show the integration of the synthetic cassettes.

**(g)** Expression of pluripotency genes was measured by RT-qPCR in WT NCRM2, Δ209, mini-1 and synHLA integration. Data represent mean ± SEM from n = 3 independent experiments. *P*-values were calculated by two-way ANOVA *t*-test (* $p<0.05$, ns = not significant).

**(h)** Representative images of the alkaline phosphatase (AP) staining for each cell line. WT NCRM2 as a control.

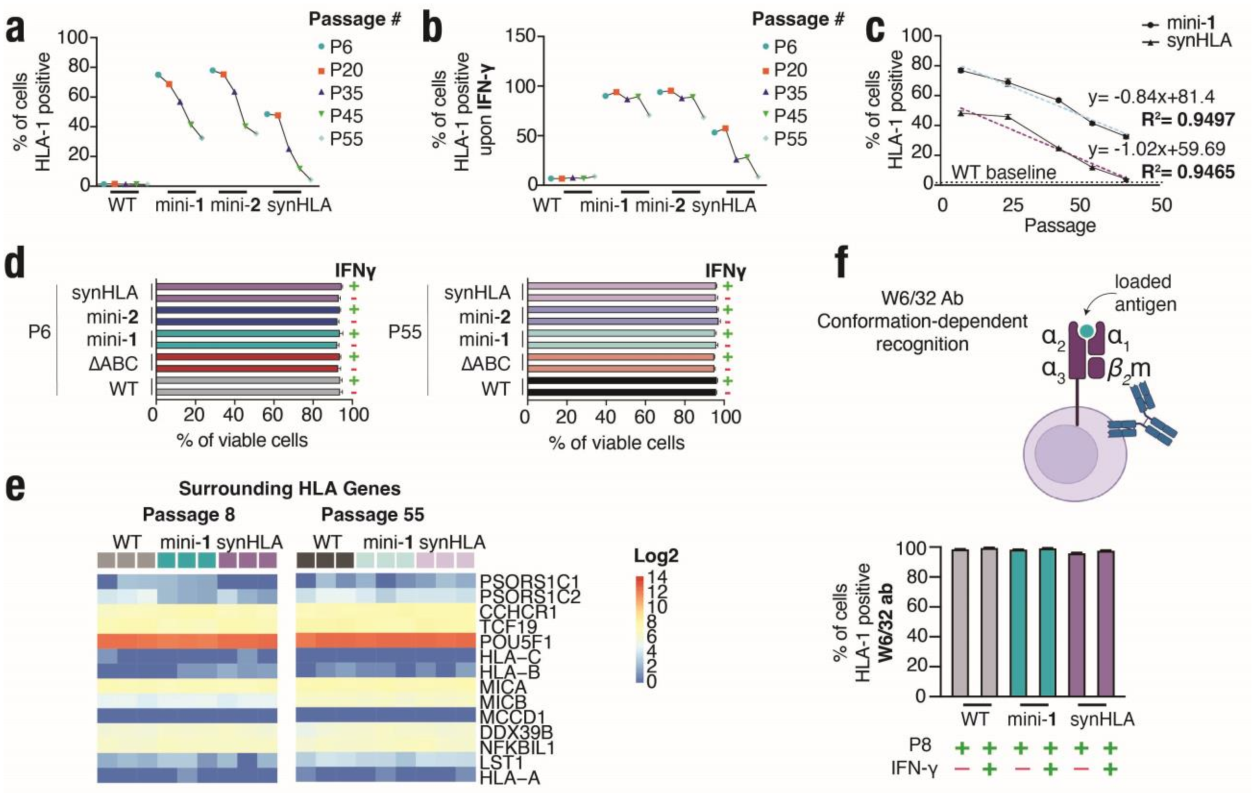


**Supplementary Figure S11. Analysis of HLA expression in engineered cells.**

**(a–b)** Class-I HLA expression was determined by flow cytometry in WT PGP1, FlpOut ΔABC, FlpOut (marker-excised) mini- and synHLA hPSCs at different passages (P6, 20, 35, 45 and 55) with and without IFN-γ induction. Class-I HLA was stained with anti-HLA class-I antibody (clone EMR8-5). Data represent mean ± SEM from n = 3 independent experiments. Two different haplotypes for the mini-synHLAs were used.

**(c)** The linear regression dashed lines show the trend of class-I HLA expression in FlpOut mini-1 (light blue dashed lines) and synHLA hPSCs (purple dashed line) over time with the corresponding slopes and coefficients of determination ($R^2$) indicated on the graph. The horizontal black dotted line indicated the WT PGP1 baseline expression. Data represent mean ± SEM from n = 3 independent experiments.**(d)** The graph shows the cell viability determined by flow cytometry in WT PGP1, FlpOut ΔABC, FlpOut mini-1 and -2, and synHLA hPSCs with and without IFN-γ induction at passage 6 (P6) and passage 55 (P55) DAPI was used as viability dye (n = 3).

**(e)** $Log_2$ gene expression heatmap of genes surrounding the HLA genes on chromosome 6 from RNA-seq analysis of WT, FlpOut (marker-excised) mini-1 and synHLA hPSCs at Passage 8 and 55.

**(f)** Cell surface class-I HLA expression was assessed by flow cytometry using the W6/32 antibody, a conformation-dependent monoclonal antibody specific for properly assembled class-I HLA heavy chain–β2-microglobulin complexes. The expression was assessed at Passage 8 (P8) in WT PGP1, FlpOut ΔABC, FlpOut (marker-excised) mini- and synHLA hPSCs with and without IFN-γ induction. Data represent mean ± SEM from n = 3 independent experiments.

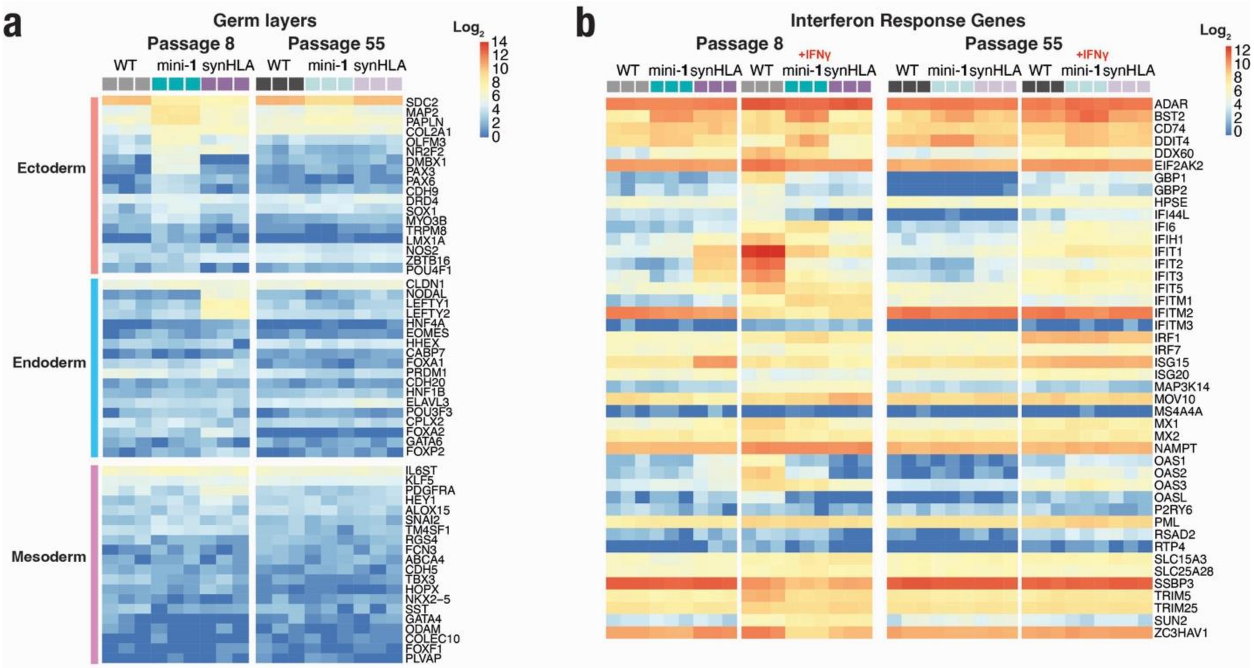


**Supplementary Fig. S12. Germ Layer and Interferon Heatmaps.**

**(a)** $Log_2$ gene expression heatmap of lineage-specific markers from RNA-seq analysis of WT, FlpOut (marker-excised) mini-1 and synHLA hPSCs at Passage 8 and 55.

**(b)** $Log_2$ gene expression heatmap of interferon response genes from RNA-seq analysis of WT, FlpOut (marker-excised) mini-1 and synHLA hPSCs at Passage 8 and 55.

**Supplementary Table S1. gRNA sequence list**

| Name | Type | [PAM] Spacer Sequence | Experiment |
|---|---|---|---|
| gRNA1 | Cas12a | [TTTC] CTCCACCATGGGCTTTGAAAA | HLA-A 5’ deletion |
| gRNA2 | Cas12a | [TTTC] TAGAGGTGCAGCTTAATTCCC | HLA-A 5’ deletion |
| gRNA3 | Cas12a | [TTTG] CCCTGTGGCTACATTAATAAA | HLA-A 3’ deletion |
| gRNA4 | Cas12a | [TTTA] TGGAACTGGTGATTCCACCA | HLA-A 3’ deletion |
| gRNA5 | Cas12a | [TTTC] CTGAAGACATATGGCTACTTG | HLA-B 5’ deletion |
| gRNA6 | Cas12a | [TTTA] GGAAGATATGAGACACTAATAAT | HLA-C 3’ deletion |
| gRNa7 | Cas9 | GGG GACTCATATAGGGAAACTCG | HLA-B 5’ deletion |
| gRNA8 | Cas9 | GGG CCATTAAGATGGTTGGGAGA | HLA-C 3’ deletion |
| gRNA9 | Cas9 | CGG GGACGTCGGAAAGTTCCGGG | HLA-DRB9 5’ deletion |
| gRNA10 | Cas9 | TGG GGTCCAACTGAGGCAATGCG | HLA-DQB1 3’ deletion |
| gRNA11 | Cas12a | [TTTG] GGATCCGCTACCCATTTCCGAGC | HLA-DRB9 5’ deletion |
| gRNA12 | Cas12a | [TTTC] ACCATGGCCACTTGCTTGTGGGA | HLA-DQB1 3’ deletion |
| gRNA13 | Cas9 | GGG GCCACTAGGGACAGGAT | LP Integration into CCR5 |
| gRNA14 | Cas9 | AGG AATATCTGTCAAAGCCAAGG | LP Integration into CD40LG |
| gRNA15 | Cas9 | TGG GCACCTCTGTAGGATCACAG | LP Integration into TRBC1 |
| gRNA16 | Cas9 | TGG TTAGTCCTAGTGCCATGAAG | LP Integration into ROGI1 |

**Supplementary Table S2. PCR primer sequence list**

| Name | Sequence | Experiment |
|---|---|---|
| CCR5 5’ junction | ACCTGCAGCTCTCATTTTCCA | LP Integration Into iPSCs |
| CCR5 3’ junction | ACAAGTCTCTCGCCTGGTTC | LP Integration Into iPSCs |
| CD40LG 5’ junction | AAGCCCAGCCTATGAATGCC | LP Integration Into iPSCs |
| CD40LG 3’ junction | GGCTGGACGGTCAGTGTTAT | LP Integration Into iPSCs |
| TRBC1 5’ junction | TTGGGCACCTGTGGTTCATT | LP Integration Into iPSCs |
| TRBC1 3’ junction | CCCCTCATCCCTCTTACGTG | LP Integration Into iPSCs |
| ROGI1 5’ junction | ATGTCTGGGTCGACGAAGTT | LP Integration Into iPSCs |
| ROGI1 3’ junction | TGGCACTACTTTTTCGCAGC | LP Integration Into iPSCs |
| ROGI2 Fw | AGCCTAACGGTGTTTAAGGAGG | |
| ROGI2 Rv | TGCTTCGGCATTCCTAGCAG | |
| HLA-B homology arm Fw | TTGGGGGTCTCTACCGCAGTGACAGAAAACAGAGTG | HLA-B deletion |
| HLA-B homology arm Rv | TAGGAGGTCTCACTTCGAAATTAAGAAAATGAGTGTGAATGT | HLA-B deletion |
| 5’ junction Fw | CTAAAGGAGAGACCAAACGA | LP Integration into B-C HLA |
| 5’ junction Rv | ATTTCAGTCGACGAAGTTCC. | LP Integration into B-C HLA |
| 3’ junction Fw | ATTGCATCGCATTGTCTGAGTAGGT | LP Integration into B-C HLA |
| 3’ junction Rv | CCCTAAGTCTGCTAAACACAGGT | LP Integration into B-C HLA |
| 5’ junction Fw | GCAGCTGCTCAGCAACAA | Hyg integration into B-C HLA |
| 5’ junction Fw | TTCTGCTTGGATCCGAAGTTCCTA | Hyg integration into B-C HLA |
| 3’ junction Fw | ATACTTCAAAAGGAATAGGAACTTC | Hyg integration into B-C HLA |

| | | |
|---|---|---|
| 3' junction Fw | TGTCTAAGCGTCCTCTAGAC | Hyg integration into B-C HLA |
| HLA-A Fw | AGGCTTTATTCACCTCAAAGTTGC | Del/Int. verification |
| HLA-A Rv | GCCACCTCTGTATAAACCCG | Del/Int. verification |
| HLA-B Fw | TTCCCCTCCTTTCCCAGAGCCA | Del/Int. verification |
| HLA-B Rv | GGGGAGGAGTGAAGAAATCCTGCA | Del/Int. verification |
| HLA-B Fw | TTCTCCATTCAACGGAGGGC | Del/Int. verification |
| HLA-B Rv | AGACTGACCGAGAGAGCCTG | Del/Int. verification |
| HLA-C Fw | GGCTGCGTGTAAGTGATGGC | Del/Int. verification |
| HLA-C Rv | AGAACATGCGGATTCTGGAAAGTT | Del/Int. verification |
| HLA-C Fw | CTGTCTCTCCACCTCCTCAC | Del/Int. verification |
| HLA-C Rv | CCATCATGGGCATCGTTGCT | Del/Int. verification |
| HLA-B/A junction Fw | GTCTCTGTGCATTCTGAGACAA | specific 38kb synHLA primer |
| HLA-B/A junction Rv | TGCTAGGGATATGACTGCTTTTG | specific 38kb synHLA primer |
| HLA-A/C junction Fw | CCAGTGAGCCAAGGATCGAAT | specific 38kb synHLA primer |
| HLA-A/C junction Rv | TGCACTGGTTAGGATGACTGTTA | specific 38kb synHLA primer |
| HLA-B/A junction Fw | AATTCTGTTCATCTTCCCTTCCAC | specific 115kb synHLA primer |
| HLA-B/A junction Rv | GATATGGGCTTTAGAATAGGGAGGT | specific 115kb synHLA primer |
| HLA-A/C junction Fw | ATGTGATAGAAGTGATTTGGACAC | specific 115kb synHLA primer |
| HLA-A/C junction Rv | CATGAATTGGAATGGGAACTCGG | specific 115kb synHLA primer |
| HLA-DRB9 homology arm Fw | TTGGGGGTCTCTACCGGGGCGATTAAGATCTCCTTCT | HLA-DRB9 deletion |
| HLA-DRB9 homology arm Rv | TAGGAGGTCTCACTTCGGAATGCAAACAGCCAGAAG | HLA-DRB9 deletion |
| HLA-DQB1 homology arm Fw | ACGAAGGTCTCGTTGTTTCCAACACACCTCTGCCTG | HLA-DQB1 deletion |
| HLA-DQB1 homology arm Rv | GGGAGGGTCTCCTAGCGTACTAATCTGCATCCCCACC | HLA-DQB1 deletion |
| 5' junction Fw | GGGATCATAAAGGGAGTGTACCA | LP Integration into DRB9-DQB1 HLA |
| 5' junction Rv | GGGGGTTATCATTGGCGTAA | LP Integration into DRB9-DQB1 HLA |
| 3' junction Fw | GGGAGGATTGGGAAGACAATAG | LP Integration into DRB9-DQB1 HLA |
| 3' junction Rv | CTGCAAGGAACATAGGAGTATAGG | LP Integration into DRB9-DQB1 HLA |
| 5' junction Fw | ATGCCAACAGCAGACTAGGT | Hyg Integration into DRB9-DQB1 HLA |
| 5' junction Fw | TTCTGCTTGGATCCGAAGTTCC | Hyg Integration into DRB9-DQB1 HLA |
| 3' junction Fw | GCTGCAACTTACCTCCGGGAT | Hyg Integration into DRB9-DQB1 HLA |
| 3' junction Fw | CCCAGGTGTATTCCTCAACCTTGGC | Hyg Integration into DRB9-DQB1 HLA |
| HLA-DRB9 Fw | CTTCCACCCGCTCTTCAACT | Del. verification |
| HLA-DRB9 Rv | AACGTGACAGAGACGCAAGT | Del. verification |

| | | |
|---|---|---|
| HLA-DRB6 Fw | GAATAATGCAATGTGTTTGTGGCA | Del. verification |
| HLA-DRB6 Rv | GGTTCGAGATCCTGGAGCAAA | Del. verification |
| HLA-DRB1 Fw | TGGTCCAATCCAGTTTCCTACC | Del. verification |
| HLA-DRB1 Rv | AGTGGACAGACTTGGCATGTATTTT | Del. verification |
| HLA-DQA1 Fw | ACAACTCTACCGCTGCTACC | Del. verification |
| HLA-DQA1 Rv | GGATATAGGAGTAAAAGGCAGGAAG | Del. verification |
| HLA-DQB1-AS1 Fw | AAGAGCGATGGACACAATGCT | Del. verification |
| HLA-DQB1-AS1 Rv | GTGATCCTTGGTCACTGGTGTT | Del. verification |
| HLA-DQB1 Fw | CATGTTTGTCCACAACTCAATTCCT | Del. verification |
| HLA-DQB1 Rv | GCCCACAATTACTCACCAAGA | Del. verification |

### Supplementary Table S3. Gene fragment list

| Name | Sequence |
|---|---|
| HLA-C 3' homology arm | ACTACAGTCGGGTCTCATTGTATGTATAAAACCAACCCATACCCCAACCACATTGAGCACATGCTCTCAGGGTCTCCTGAGGGACGTGTCATGGGCTGTGGTCACTCGTATTTGGCTCAGAATAAATCTCTTCAAATATTTTATGAAGTTTGCCTCTTTTCATTGACAGTATGGAATTCTGATGTAGTAAGAGGGTTCAAGTGCTGGAGTGTGAAGGGTGGGAAAAGAATGATAAATTTTAATTATTGGAGCAGTGCTCCAAGACAAGAAATTTATCTAGTATCTGGTAGGGATTCAGAGTGTCTGAATGAACCGGTGACTAATAAATAACATCTTTCCACCCATTCCCTTGAAAATAAGTTATTACATCAAGTTTTTGTCTATCCCAGTTCATACTCCAGATTATTGGAGTGGCATGGTGTGCTTGGCATTGTCTCTTACTGGGATTAAAACACCCAGGTTTGTAAGATCGCTATGAGACCCACGATTAGC |

### Supplementary Table S4. qPCR primer list

| Name | Sequence | Usage |
|---|---|---|
| Dnmt3b Fw | GACTCGATCCTCGTCAACGG | pluripotency |
| Dnmt3b Rv | ATCTTCGGCCTCTGATCTCCG | pluripotency |
| Nanog Fw | GAGATGCCTCACACGGAGAC | pluripotency |
| Nanog Rv | GGGTTGTTTGCCTTTGGGAC | pluripotency |
| Oct4 Fw | AGTGAGAGGCAACCTGGAGA | pluripotency |
| Oct4 Rv | ACACTCGGACCACATCCTTC | pluripotency |
| Sox2 Fw | GCACAACTCGGAGATCAG | pluripotency |
| Sox2 Rv | CAGCGTGTACTTATCCTTCT | pluripotency |
| Klf4 Fw | AGCCTAAATGATGGTGCTTGGT | pluripotency |
| Klf4 Rv | CCTTGTCAAAGTATGCAGCAGT | pluripotency |
| RSP29 Fw | AATATGTGCCGCCAGTGTTT | pluripotency |
| RSP29 Rv | CCCGGATAATCCTCTGAAGG | pluripotency |
| FOXN1 Fw | CAAAACATGCCAGCGCCGCTTT | TEC |
| FOXN1 Rv | GCCTGGTGCAATGGACTGCCTT | TEC |
| DLL4 Fw | CGGGTACCTTCTCGCTCATC | TEC |
| DLL4 Rv | ATGAGTGCATCTGGTGGCAA | TEC |
| Puromycin Fw | GACATCGGCAAGGTGTGGGTCG | Marker cassette |
| Puromycin Rv | GGAACCGCTCAACTCGGCCATG | Marker cassette |
| GFP Fw | AACATGGCCTCTCTCCCAGcgA | Marker cassette |

| | | |
|---|---|---|
| GFP Rv | TTTGGATTGCCGGTGCCCTGAC | Marker cassette |
| Hygromycin Fw | AAAGCCTGAACTCACCGCGACG | Marker cassette |
| Hygromycin Rv | GCACGAGATTCTTCGCCCTCCG | Marker cassette |

### Supplementary Table S5. Antibodies List

| Name | Company | Dilution |
|---|---|---|
| Anti-HLA Class1 ABC antibody, mouse monoclonal | Abcam [EMR8-5] | 1:50 (flow cytometry) |
| Anti-HLA Class1 ABC antibody, mouse monoclonal | Invitrogen [W6/32] | 1:100 (flow cytometry) |
| Recombinant Anti-CD31/PECAM-1 Antibody, Rabbit Monoclonal | Sino Biological [10148-R078] | 1:100 (IHC) |
| Polyclonal goat IgG Human Oct3/4 antibody | R&D Systems [AF1997] | 1:200 (IHC) |
| Polyclonal goat IgG Human Nanog antibody | R&D Systems [AF1759] | 1:200 (IHC) |
| Goat anti-Rabbit IgG secondary antibody, Alexa Fluor™ Plus 647 | Invitrogen [A32733] | 1:1000 |
| Goat anti-Mouse IgG secondary antibody, Alexa Fluor™ 647 | Invitrogen [A21236] | 1:1000 (IHC), 1:500 (flow cytometry) |
| Donkey anti-Goat IgG Secondary Antibody, Alexa Fluor™ 647 | Invitrogen [A21447] | 1:1000 |
| DAPI | BD Pharmingen [564907] | 1:2000 (flow cytometry) |
